# Isoform resolved measurements of absolute translational efficiency define interplay of HIF and mTOR dysregulation in kidney cancer

**DOI:** 10.1101/2021.07.06.451052

**Authors:** Yoichiro Sugimoto, Peter J. Ratcliffe

**Affiliations:** The Francis Crick Institute, 1 Midland Road, London NW1 1AT, UK; Ludwig Institute for Cancer Research, Nuffield Department of Clinical Medicine, University of Oxford, Oxford OX3 7FZ, UK

## Abstract

Hypoxia inducible factor (HIF) and mammalian target of rapamycin (mTOR) pathways orchestrate responses to oxygen and nutrient availability. These pathways are frequently dysregulated in cancer, but their interplay is poorly understood, in part because of difficulties in simultaneous measurement of global and mRNA-specific translation. Here we describe a workflow for measurement of absolute translational efficiency of mRNAs resolved by their transcription start sites (TSSs). Its application to kidney cancer cells revealed a remarkable extent of translational reprogramming by mTOR, strongly affecting many metabolic enzymes and pathways. By contrast, global effects of HIF on translation were limited, and we did not observe reported translational activation by HIF2A. In contrast, HIF-dependent alterations in TSS usage were associated with robust changes in translational efficiency in a subset of genes. Analyses of the interplay of HIF and mTOR revealed that specific classes of HIF1A and HIF2A transcriptional target gene manifest markedly different sensitivity to mTOR, in a manner that supports combined use of HIF2A and mTOR inhibitors in treatment of kidney cancer.

## Introduction

Precise regulation of transcription and translation is required to define patterns of protein synthesis in healthy cells. Nevertheless, attempts to understand disease have often focused on a single pathway of transcriptional or translational control, despite simultaneous dysregulation of these processes. For instance, two major pathways that link the cellular environment to gene expression, the HIF and mTOR pathways, are both dysregulated in many types of cancer. The most common kidney cancer, clear cell renal carcinoma, is characterized by constitutive upregulation of the HIF pathway due to defective function of its conditional E3 ubiquitin ligase, the von Hippel-Lindau tumour suppressor gene (*VHL*), and by hyperactivation of mTOR^1, 2^. On the other hand, oxygen depletion, or hypoxia, often created in the tumour environment, increases the activity of HIF and inhibits mTOR pathways^3, 4^.

HIF mediates the cellular response to hypoxia through a well-defined role in transcription, but recent studies also report a role in translation^3^. In the presence of oxygen, two isoforms of HIFα proteins (HIF1A and HIF2A) are ubiquitinated by VHL and degraded^3^. This prevents the formation of transcriptionally active heterodimers with HIF1B. In addition, HIF2A is reported to regulate translation via a HIF2A-specific mode of non-canonical cap-dependent translation that is mediated by eukaryotic translation initiation factor 4E family member 2 (EIF4E2)^5^. It was further reported that a large subset of genes, including HIF transcription targets, are translationally upregulated by the HIF2A/EIF4E2 axis resulting in induction of protein in hypoxic cells even when HIF dependent transcription was ablated by HIF1B knockdown^6^. Evaluation of this mode of action of HIF is important given recent efforts to treat *VHL*-defective kidney cancer through HIF2A-HIF1B dimerization inhibitors^7, 8^ whose action to prevent transcription might be circumvented by this activity of HIF2A on translation. Apart from this, it is generally unclear how HIF dependent transcription interacts with the regulation of translation.

mTOR forms two different complexes, mTORC1 and mTORC2. mTORC1 dependent control of translation via EIF4E binding protein (EIF4EBP) phosphorylation is a key regulator of responses to nutrient availability, cellular stress and growth factors^9, 10^. When mTORC1 is inhibited, such as by nutrient deprivation, unphosphorylated EIF4EBP binds to EIF4E and this blocks the EIF4E-EIF4G1 interaction, which is necessary to form a canonical translation initiation complex^10^. On the other hand, mTORC2, which is stimulated by insulin, controls cell proliferation and migration by phosphorylating AKT serine/threonine kinase and other targets^9^. Comprehensive characterization of the regulation of gene expression by the HIF/VHL and mTOR pathways is crucial to understanding the biology of *VHL*-defective kidney cancer, particularly as agents targeting both these pathways are being deployed therapeutically^11, 12^. Although mTOR has been reported to be inhibited by HIF under hypoxia^4^, its interactions with the HIF system are poorly understood. In particular, the interactions between HIF-dependent and mTOR-dependent transcriptional and translational targets across the genome have not been defined.

In part, this reflects the lack of highly efficient methods to measure translational efficiency and to interface such measurements with transcriptional data. Ideally, such a method would measure absolute translational efficiency at the level of the specific transcript. Direct measurement of absolute translational efficiency greatly facilitates comparison across different conditions since alterations in global translational efficiency may otherwise confound measures of relative translational efficiency^13^. Resolution of specific transcripts by their TSS provides important insights into the mode of translational regulation and is particularly important when assessing translation in the setting of a large transcriptional change as often occurs in cancer^14, 15^.

Here we describe the development and validation of a new method, high-resolution polysome profiling followed by sequencing of the 5 ′ ends of mRNAs (HP5), that addresses these challenges, and demonstrate its use in defining the interplay between transcriptional and translational regulation by the HIF/VHL and mTOR signalling pathways in *VHL*-defective kidney cancer cells.

## Results

### Establishment of HP5 workflow

HP5 encompasses five key features (Fig. 1a and Supplementary Fig. 1). First, cell lysate is fractionated on a sucrose gradient to resolve mRNAs down to the associated ribosome number (from 1 to 8 or more). Second, an equal amount of an external RNA standard is added to each fraction for the precise quantification of mRNA abundance. Third, cDNA libraries, indexed with respect to the fraction and experimental sample are prepared from the extracted total RNA without purification steps or tube/plate transfers. The fractionated samples are then multiplexed, enabling parallel processing at scale. Up to 16 samples were multiplexed in this study, but the method is amenable to a larger number of samples, as permitted by the diversity of the indexing sequence. Fourth, 3′ ends of cDNAs (i.e. 5′ ends of mRNAs) are converted into libraries compatible with high-throughput DNA sequencing utilizing Tn5 transposase. Finally, mean ribosome load of mRNA isoforms^16^, resolved by their TSS, is calculated from the sequencing data.

**Fig. 1.**
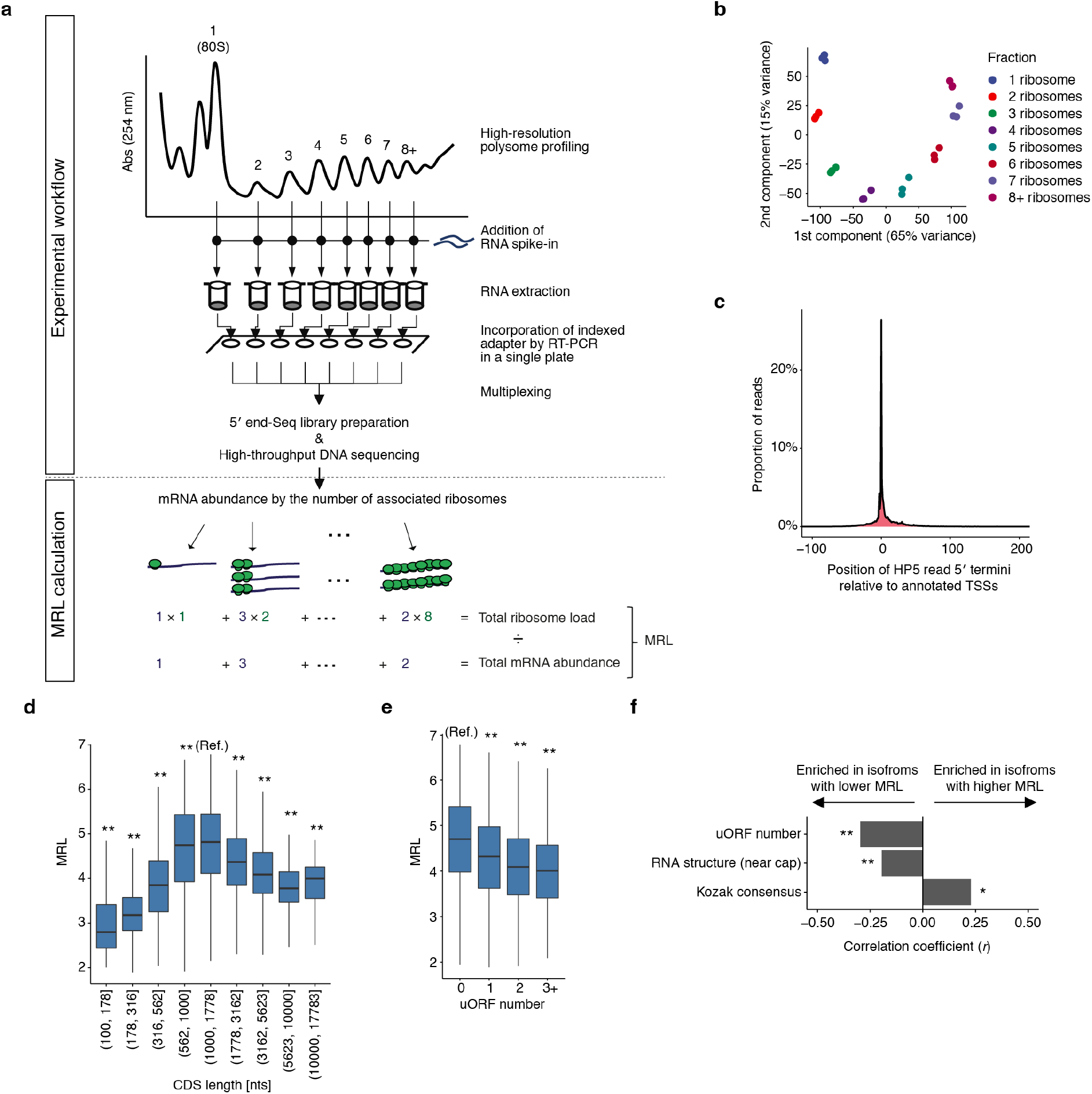
HP5 reliably measured translational efficiency of mRNAs resolved by their TSS. **(a)** Schematic overview of HP5 strategy. Top panel shows the experimental workflow including polysome profile, whereas bottom panel shows an example of the mean ribosome load (MRL) calculation. **(b)** Principal component analysis of HP5 data by polysome fraction, for 3 independent RCC4 VHL clones. **(c)** Exact position of identified 5′ termini relative to the closest annotated TSSs (RefSeq and GENCODE); data is the proportion of HP5 reads with the indicated 5′ terminus, relative to the total reads mapping to that gene locus **(d)** MRL as a function of CDS length. The distribution of MRL for mRNAs with the indicated CDS length was compared to that for a CDS length of 1,000 to 1,778 nucleotides (nts), using the Mann-Whitney *U* test. **(e)** MRL as a function of uORF number. The distribution of MRL for mRNAs with the indicated uORF number was compared to that without an uORF, using the Mann-Whitney *U* test. **(f)** mRNA features associated with higher or lower MRL in the two most differentially translated mRNA isoforms (FDR < 0.1) derived from same gene, but differing by their TSS. Comparisons were performed for those pairs of the mRNA isoforms which differed in the relevant features; the effect size of the association with each mRNA feature was measured using matched-pairs rank biserial correlation coefficients and significance determined by the Wilcoxon signed-rank test. RNA structure (near cap), is the stability of predicted RNA secondary structures (expressed as the inverse of the minimum free energy per nucleotide, first 75nts of mRNAs); Kozak consensus, match score to the consensus Kozak sequence. *: *p* < 0.05, ** *p* < 0.005. *P* values were adjusted for multiple comparisons using Holm’s method. Details of the sample sizes and exact *p* values for (d-f) are summarized in Supplementary Notes.

We first evaluated the basic performance of HP5 using RCC4 VHL cells, in which constitutive upregulation of the HIF system in *VHL*-defective RCC4 cells has been restored to normal by stable transfection of an intact *VHL* gene (Supplementary Fig. 2). We obtained an average of 3.3 million reads per fraction with ∼80% of reads mapping to mRNA (Supplementary Table 1). Importantly, by the exclusion of mRNA or cDNA purification steps before the first PCR amplification and multiplexing of samples at an early stage of the protocol, HP5 successfully generated each library from 100-fold less total RNA compared to a similar method (∼30 ng compared to 3 µg)^17^, enabling efficient analysis of large numbers of samples and/or materials with low RNA yield. HP5 was highly reproducible, principal component analysis of mRNA abundance data demonstrating tight clustering by the polysome fraction, across three clones of RCC4 VHL cells (Fig. 1b). Furthermore, the 5′ terminus of HP5 reads matched annotated TSSs in RefSeq or GENCODE precisely at nucleotide resolution, confirming the accuracy of 5′ terminal mapping (Fig. 1c).

To test the performance of HP5 against previous work in the field, we examined the overall relationships between the translational efficiency and selected mRNA features including those with known associations with translational control. Translational efficiency was calculated as the mean ribosomal load for each of 12,459 mRNA isoforms resolved by their TSS from 7,815 genes. Using a generalized additive model, we identified varying predictive power of different mRNA features (Supplementary Fig. 3a). The four most predictive features together explained around 36% variance in mean ribosome load between mRNAs. Interestingly, coding sequence (CDS) length showed the clearest association with mean ribosome load, values being greatest for mRNAs with a CDS length of around 1,000 nucleotides, but declining progressively with longer CDS (Fig. 1d) probably due to the lower likelihood of promoting re-initiation of translation by mRNA circularization^18^. In agreement with previous studies^16, 19, 20^, analysis of HP5 data identified the negative effect on translation of upstream open reading frames (uORFs) and RNA structures near the cap, as well as the positive effect of the Kozak consensus sequence (Fig. 1e and Supplementary Fig. 3b and c). Importantly, the association of mean ribosome load with mRNA features known to affect translation also extended to comparisons between mRNA isoforms derived from the same gene by alternate TSS usage (Fig. 1f). Overall therefore, HP5 reproduced and extended known associations between mRNA features and translation, verifying its performance in the measurement of translational efficiency at transcript resolution.

### HP5 revealed that the extent of mTOR dependent translation regulation is greater than reported

We next applied HP5 to the analysis of mTOR pathways, which are frequently dysregulated along with hypoxia signalling pathways in *VHL*-defective kidney cancer. To analyse translational changes that are directly consequent on mTOR inhibition, RCC4 VHL cells were treated for a short period (2 hours) with Torin 1, an ATP competitive inhibitor of mTORC1 and mTORC2^21^, before analysis by HP5. mTOR inhibition globally suppressed translation as shown by a marked reduction in polysome abundance (Fig. 2a). Measurements of absolute changes in translational efficiency were performed by calculation of mean ribosomal load using the external standards, and initially analysed at the level of the gene. This provided the first direct display of both general reduction in translation by mTOR inhibition and of its heterogeneous effects on individual genes across the genome (Fig. 2b). mTOR has been reported to regulate a wide range of processes by different mechanisms^9^, while its directly identified translational targets have been more limited, particularly involving proteins that function in translation itself. Our data confirmed many of these known mTOR translational targets, as well as the previously described resistance of many transcription factors^10^. Importantly, it also demonstrated directly that the translation of genes encoding proteins with many other functions, such as in different metabolic pathways, and in proteasomal degradation is also hypersensitive to mTOR inhibition (Fig. 2c).

**Fig. 2.**
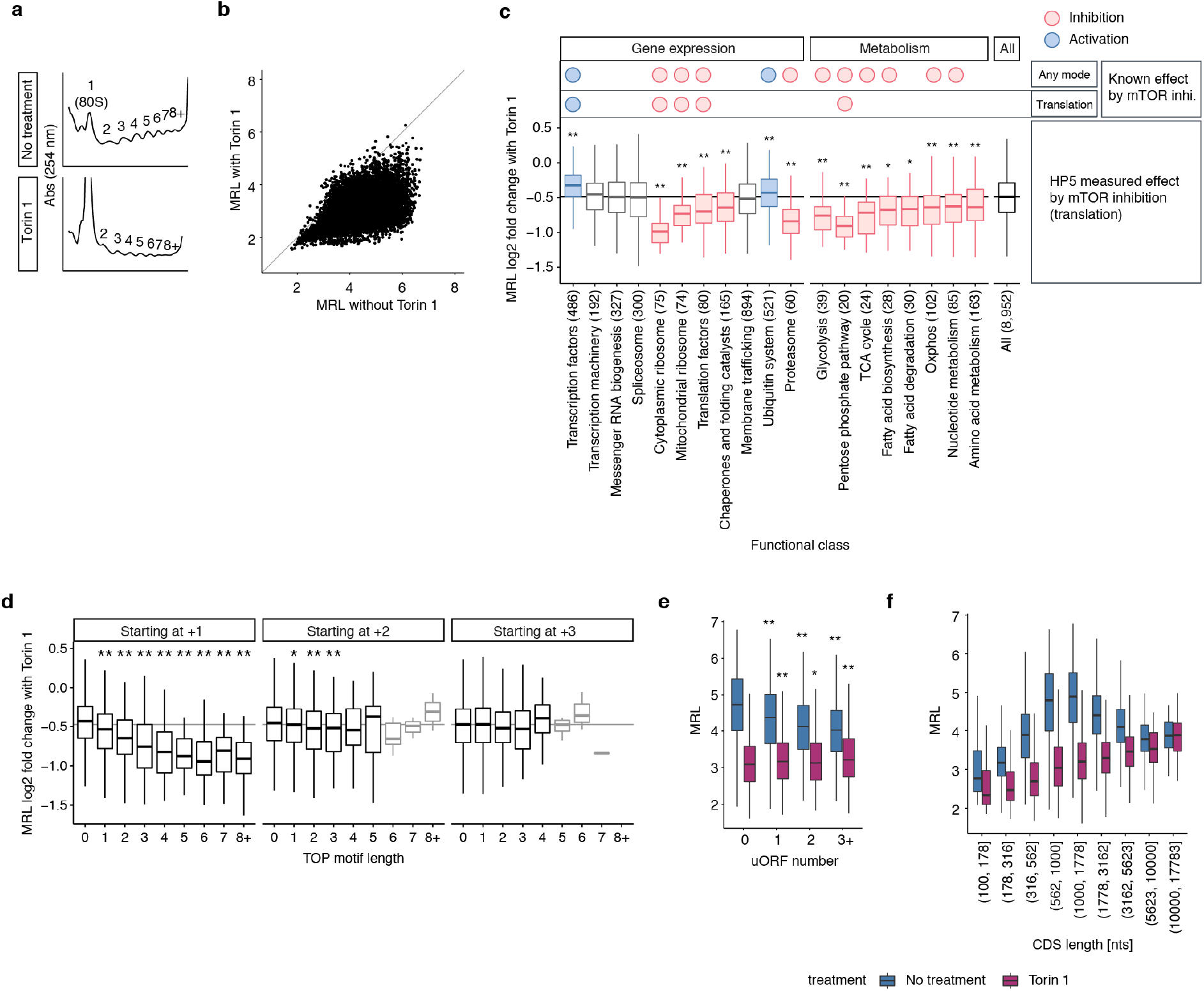
Comprehensive analysis of mTOR dependent translation regulation by HP5. **(a)** Polysome profiles of RCC4 VHL cells with and without Torin 1 treatment (250 nM, 2 hours). **(b)** Comparison of the mean ribosome load (MRL) of genes with and without Torin 1 treatment. Each point represents the average of data for independent clones of RCC4 VHL cells; n = 2 (with Torin 1), n = 3 (without Torin 1) **(c)** Box plots showing changes in translational efficiency of genes (expressed as log2 fold change MRL) after exposure to Torin 1, among different functional classes as defined by KEGG orthology. Known systematic regulation of each functional class by mTOR inhibition by any mechanism or specifically by translational regulation, is indicated above the boxplots (see Methods for definitions). Responses within a functional class were compared against responses for all other genes using the Mann-Whitney *U* test; classes more downregulated or less downregulated compared to all other genes (i.e. hypersensitive or resistant to mTOR inhibition, *p* < 0.05) are colored red or blue respectively; the number of genes in each class is indicated in parenthesis. **(d)** Box plots showing changes in translational efficiency upon exposure to Torin 1 as a function of TOP motif (pyrimidine tract) length (x-axis) and starting position with respect to the mRNA cap (individual panels). The distribution of MRL for mRNAs with the indicated TOP motif length was compared to that without a TOP motif using the Mann-Whitney *U* test; boxes representing less than 10 mRNAs are faded. **(e-f)** Box plots showing MRL as a function of **(e)** uORF number, **(f)** CDS length, in the presence (purple) or absence (blue) of Torin 1. For **(e)**, the distribution of MRL for mRNAs with the indicated uORFs number was compared to that of those without a uORF using the Mann-Whitney *U* test. *: *p* < 0.05, ** *p* < 0.005. *P* values were adjusted for multiple comparisons using Holm’s method. Details of the sample sizes and exact *p* values for (c-f) are summarized in Supplementary Notes.

Direct genome-wide evaluation of translational changes produced by mTOR inhibition has not been performed on mRNAs resolved by TSS usage. The resolution provided by HP5 therefore provided an opportunity to improve the understanding of transcript-specific mRNA features associated with mTOR hypersensitivity or resistance. mTOR has been shown to regulate mRNAs with a 5′ terminal oligopyrimidine (TOP) motif. This process is reported to operate in a tract length dependent manner via the binding of La-related protein 1 (LARP1)^20^. In keeping with this, we observed that mRNAs with longer TOP motifs commencing immediately after the cap were more strongly downregulated upon mTOR inhibition (Fig. 2d). In contrast, although it has been reported that TOP motifs starting between +2 and +4 nucleotides downstream of the cap also mediate mTOR control^10^, the high-resolution analysis permitted by HP5 revealed that any such association with Torin 1 sensitivity was very much weaker if the TOP motifs did not start immediately after the cap (Fig. 2d). Thus, precise positioning of the cap and TOP motif appears to be critical.

Although this data confirmed the importance of the TOP motif for translational regulation by mTOR, the proportion of mRNAs containing a TOP motif was limited (only 6% of mRNAs had TOP motif greater than 2 nucleotides, Supplementary Fig. 4a) compared to the global extent of translational alteration by mTOR inhibition, implying that additional mechanisms determine the mTOR sensitivity. To explore this, we examined the interaction of Torin 1 induced changes in translation with uORF frequency and CDS length, the two most important mRNA features affecting translational efficiency under mTOR active conditions (Supplementary Fig. 3a). Strikingly, we observed that uORF number retained only a very weak association with mean ribosome load under mTOR inhibition (Fig. 2e). In respect of CDS length, the increased translational efficiency of mRNAs with near 1 kb CDS was not observed upon mTOR inhibition (Fig. 2f and Supplementary Fig. 4b). Rather, there was a progressive increase in mean ribosomal load with increasing CDS length, as might be expected if CDS length was not affecting translational initiation. These differences suggest that mTOR pathways also impinge on the translational effects of these mRNA features. For instance, EIF4EBP activation by mTOR inhibition might prevent mRNAs from forming a loop through blocking EIF4E and EIF4G1 interactions. Taken together, our findings demonstrated that HP5 can measure absolute translation change at a TSS isoform resolution with high sensitivity. Overall the analyses revealed that the extent of translation regulation by mTOR is greater than previously reported and have refined the understanding of mRNA features that influence mTOR sensitivity.

### Limited role of HIF2A in regulating translation compared to HIF1A

We next sought to examine translational regulation by HIF/VHL pathway by applying HP5 to *VHL*-defective RCC4 and 786-O cells re-expressing either wild type VHL (RCC4 VHL and 786-O VHL) or empty vector alone. The two cell lines were chosen because RCC4 expresses both HIF1A and HIF2A whereas 786-O expresses only HIF2A (Supplementary Fig. 2), enabling us to distinguish between the roles of HIF1A and HIF2A. Furthermore, previous studies reporting the role of HIF2A/EIF4E2 pathway were performed in part using 786-O cells^5, 6^. Figure 3a shows the changes in translational efficiency associated with loss of *VHL* for RCC4 cells (upper panel), or 786-O cells (middle panel) compared to the action of Torin 1 on RCC4 VHL cells (lower panel). In both RCC4 and 786-O cells, *VHL*-defective status was associated with a small global downregulation of translation, with a greater number of genes showing reduced translational efficiency in *VHL*-defective RCC4 cells.

**Fig. 3.**
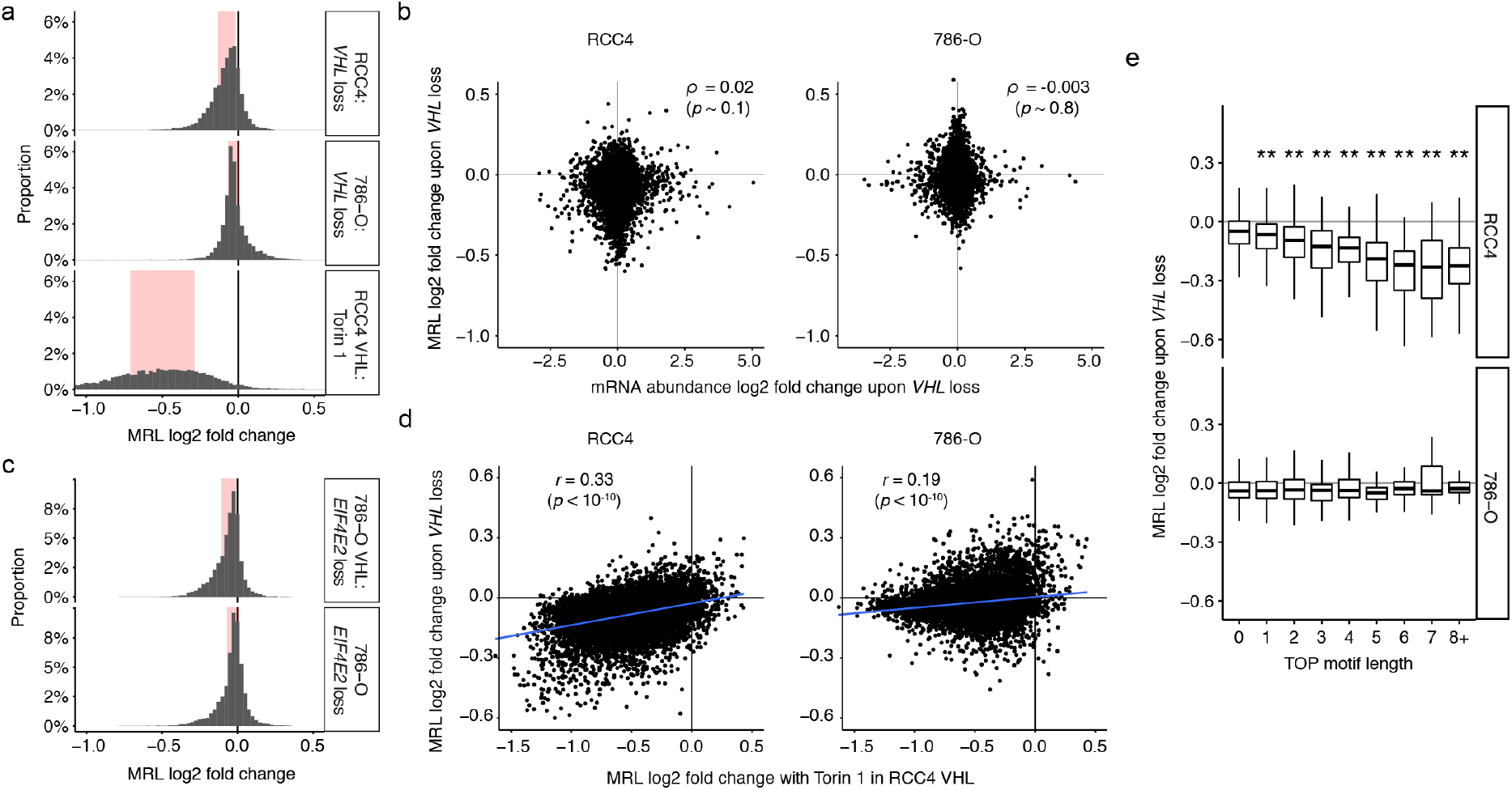
Global view of HIF dependent translational regulation. **(a)** Frequency histograms comparing changes in translational efficiency, expressed as log2 fold change in mean ribosome load (MRL) for each gene, for three interventions; *VHL* loss in RCC4 cells (upper panel), *VHL* loss in 786-O cells (middle panel), and Torin 1 treatment in RCC4 VHL cells (lower panel). Interquartile range is highlighted in red; the width of histogram bin was set to 0.05. **(b)** Scatter plots comparing changes in mRNA abundance of genes induced by *VHL* loss with the changes in translational efficiency in the respective cell type. Spearman’s rank-order correlation coefficient was used to assess the association (n = 9,318 and 7,844 for RCC4 and 786-O respectively). **(c)** Frequency histograms as in **a,** showing effects of *EIF4E2* inactivation in 786-O VHL cells (upper panel) and 786-O cells (lower panel). **(d)** Scatter plots comparing changes in translational efficiency of genes upon *VHL* loss in RCC4 (left panel) or 786-O (right panel) cells with those induced by Torin 1 treatment in RCC4 VHL cells. The blue line indicates the linear model fit by ordinary least squares. Pearson’s product moment correlation coefficient was used to assess the association (n = 8,829 and 7,512 for RCC4 and 786-O respectively). **(e)** Box plots showing changes in translational efficiency of mRNAs upon *VHL* loss as a function of TOP motif length (x-axis). Only TOP motifs starting immediately after cap were analyzed. The distribution of MRL for mRNAs with indicated TOP motif length was compared to that without a TOP motif using the Mann-Whitney *U* test. * *p* < 0.05, ** *p* < 0.005. *P* values were adjusted for multiple comparisons using Holm’s method. Details of the sample sizes and exact *p* values are summarized in Supplementary Notes.

Particular striking, in view of the reported role of HIF2A in translational up-regulation^5, 6^, was the absence of clear up-regulation in translational efficiency in *VHL*-defective RCC4 and 786-O cells, both of which manifest strong induction of HIF2A (Supplementary Fig. 2). It is possible that HIF2A upregulates the translation of only a small number of mRNAs, for instance a subset of HIF induced mRNAs. We therefore compared changes in mRNA abundance induced by *VHL* with changes in translational efficiency or absolute translational efficiency, in the cells lacking VHL. However, we saw no correlation between regulation of transcript abundance and translation, as might have been anticipated if a set of HIF transcriptional targets were also regulated by altered translational efficiency (Spearman’s *ρ* = 0.02 and −0.003, *p* = 0.1 and 0.8 for changes in translational efficiency against changes in mRNA abundance in RCC4 and 786-O cells respectively; Fig. 3b and Supplementary Fig. 5a).

Since the action of HIF2A to promote translation has been proposed to be mediated by the non-canonical cap binding initiation factor EIF4E2^5^, we engineered *EIF4E2*-defective 786-O and 786-O VHL cells by CRISPR/Cas9 mediated inactivation, and examined effects on translational efficiency using HP5. In both 786-O and 786-O VHL cells, *EIF4E2* inactivation weakly but globally downregulated the translational efficiency of genes (Fig. 3c). If co-operation of EIF4E2 and HIF2A had a major role in translation it would be predicted that *EIF4E2* inactivation would have a larger effect in the presence of HIF2A than in its absence and hence that effects of *EIF4E2* inactivation would be substantially greater in the absence of VHL. However, we observed no evidence of this (Fig. 3c, compare upper and lower panels). Finally, to exclude the possibility that HP5 analysis did not capture the effect of HIF2A/EIF4E2 dependent translational regulation, we examined changes in protein abundance of previously reported HIF2A/EIF4E2 target genes^5, 6^ as a function of *VHL* or *EIF4E2* status in 786-O cells, using immunoblotting. This further confirmed that the effect of HIF2A/EIF4E2 pathway was considerably weaker or undetectable compared to that of HIF2A/VHL dependent transcriptional regulation (Supplementary Fig. 5b). Taken together, the data revealed little or no role for the HIF2A/EIF4E2 axis in regulation of translation under the conditions that we analysed.

Whilst we did not observe systematic upregulation of translational efficiency either of HIF transcriptional targets or other genes in *VHL*-defective cells, we did observe downregulation of translational efficiency, particularly in RCC4 cells. To examine whether this might reflect interaction of HIF and mTOR pathways, we first compared the gene-specific effects on translation that are associated with *VHL*-defective status in RCC4 cells with those observed by inhibition of mTOR with Torin 1 in RCC4 VHL cells. This revealed a moderate, but highly significant, correlation between responses to the two interventions in RCC4 cells (Pearson’s *r* = 0.33, *p* < 10^-10^, Fig. 3d left panel), Furthermore, when VHL-dependent changes in translational efficiency were related to the length of the TOP motif, mRNAs with longer TOP motif were found to be more strongly repressed by *VHL* loss in RCC4 cells (Fig. 3e upper panel). In earlier work it has been suggested that induction of HIFα, particular the HIF1A isoform can suppress mTOR pathways^4, 22^. Consistent with this, we observed that *VHL* loss in RCC4 cells was associated with a significant upregulation of mRNAs that encode negative regulators of mTOR (BNIP3 and DDIT4) or its target, the translational repressor EIF4EBP1 (Supplementary Fig. 5c). In contrast, in 786-O cells which do not express HIF1A, we observed less downregulation of translation by *VHL* loss, less association of any gene-specific effects with mTOR targets (defined either by responsiveness to Torin 1, or the length of the TOP sequence) and weaker regulation by VHL of mRNAs that repress mTOR pathways (Fig. 3d right panel and Fig. 3e lower panel, and Supplementary Fig. 5c). Taken together the findings suggest that downregulation of translation occurs in RCC4 cells as a consequence of HIF1A dependent actions on mTOR pathways, but that these effects are modest by comparison with pharmacological mTOR inhibition.

### HIF promotes alternate TSS usage to regulate translation

Although transcription may regulate translation by promoting alterative TSS usage and altering the regulatory features of the mRNA, neither HIF-dependent alternate transcriptional initiation (TSS usage) nor its effect on translation have been studied systematically. To address this, we first compared 5′ end-Seq reads from total mRNAs (i.e. unfractionated RNA) in RCC4 VHL versus RCC4 and identified 149 genes that manifest a VHL-dependent change in TSS usage (FDR < 0.1). For these genes, we defined a VHL-dependent alternate TSS (that showing the largest change in mRNA abundance with *VHL* loss). Discordant regulation of the alternate and the other TSSs (i.e. up versus down) was rare (9/149); following *VHL* loss the alternate TSS was induced in 85 genes and repressed in 64 genes (Supplementary Table 2). To test the generality of these findings and consider the mechanism, we performed similar analyses of alternative TSS usage amongst these 149 genes in sets of related conditions and compared the results (Supplementary Fig. 6). A strong correlation (Pearson’s *r* = 0.60, *p* < 10^-10^) was observed with alternate TSS usage in 786-O VHL versus 786-O cells. In contrast there was no correlation with the alternate TSS usage in 786-O VHL versus 786-O cells in which HIF transcription had been ablated by CRISPR/Cas9-mediated inactivation of *HIF1B* (Pearson’s *r* = −0.01, *p* ∼ 0.9) indicating that the effects on alternate TSS usage were dependent on HIF. In keeping with this a very striking correlation was observed between changes mediated by loss of *VHL* in RCC4 and those induced by hypoxia in RCC4 VHL cells (Pearson’s *r* = 0.85, *p* < 10^-10^). Taken together these findings reveal robust changes in TSS usage when HIF is activated.

Comparison of alternate TSS mRNA isoforms with other mRNA isoforms of same gene revealed that 64% of alternate TSS mRNA isoforms significantly differed in their polysome distribution (FDR < 0.1, Supplementary Fig. 7 and Supplementary Table 2). We therefore sought to determine to what extent the alternative TSS usage amongst these genes accounted for overall changes in their translational efficiency. To assess this, we recalculated changes in translational efficiency for each gene, omitting either (i) the VHL-dependent changes in translational efficiency within each mRNA isoform, or (ii) the VHL-dependent changes in TSS usage, from the calculation and compared the results with the experimental measurement, as derived from both parameters. The correlation was much stronger using (i) than (ii) (Pearson’s *r* = 0.83 and *r* = 0.54, *p* < 10^-10^ and 10^-5^ respectively, Fig. 4a) indicating that VHL-dependent alternate TSS usage was the primary mode of the translational regulation for this set of genes.

**Fig. 4.**
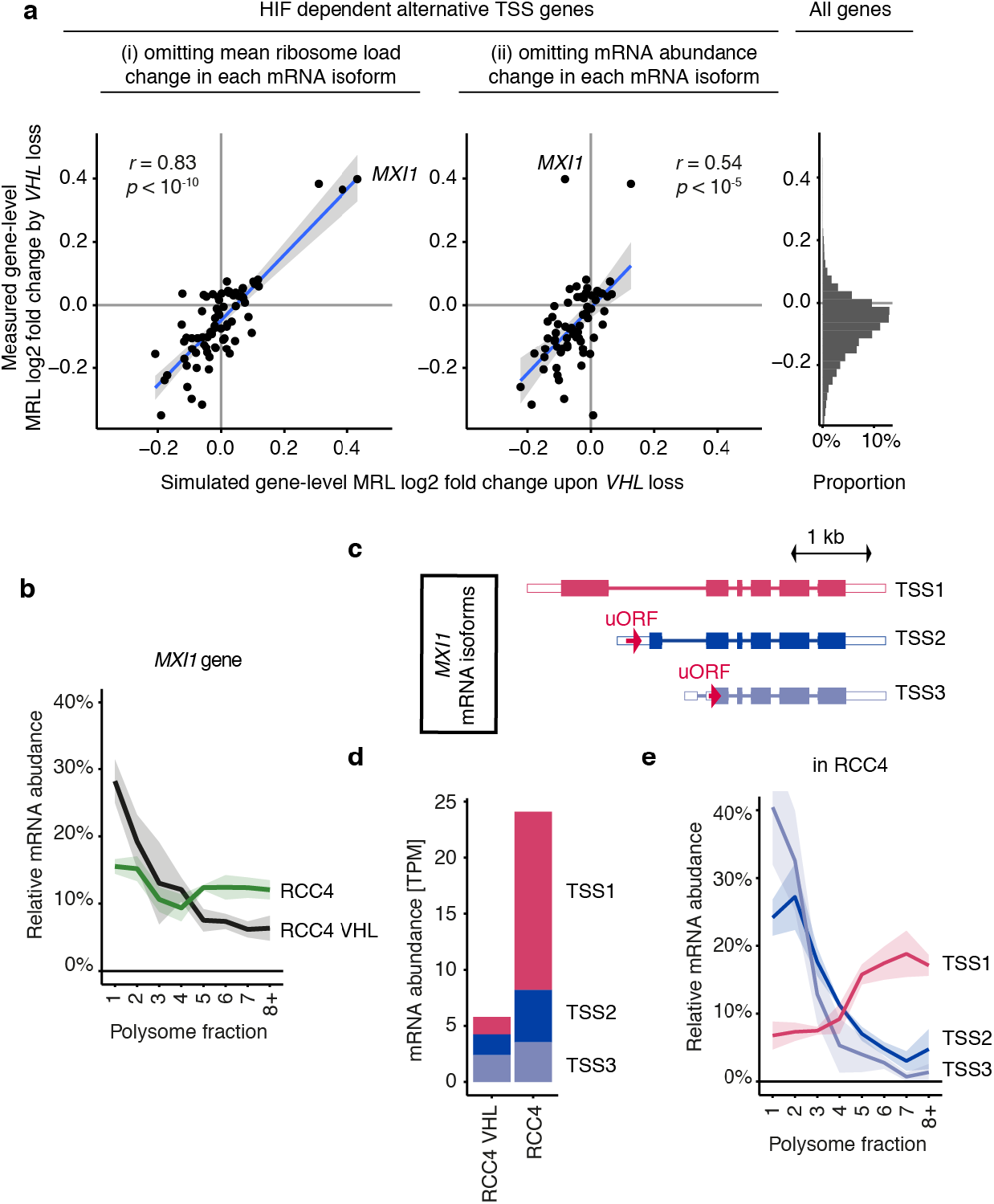
Translational regulation by VHL-dependent alternative TSS usage. **(a)** Contribution of VHL-dependent alternative TSS usage to changes in translational efficiency of genes following *VHL* loss in RCC4 cells. The scatter plots show the correlations between measured changes in translational efficiency (log2 fold change in overall mean ribosome load (MRL) for each gene, y-axis) and that simulated when omitting one parameter (x-axis). The analyses are of those genes manifesting an altered polysome distribution on their VHL-dependent alternative transcript. It demonstrates that it is this property, rather than VHL-dependant changes in mean ribosome load within transcripts, that makes the greatest contribution to overall VHL-dependent changes in translational efficiency of the gene. Pearson’s product moment correlation coefficient was used to assess the association (n = 75 and 70 for (i) and (ii), respectively). The blue line indicates the linear model fit by ordinary least squares and the grey shade shows the standard error. Right panel (the same data as in the upper panel of Fig. 3a), is provided to reference the distribution of changes in translational efficiency amongst the subset of genes manifesting alternative TSS usage to all expressed genes. **(b)** Proportion of *MXI1* mRNA distributed across polysome fractions; shaded area shows the standard deviation of the data from the three independent clones. **(c)** Schematic diagrams for the 3 the most abundant mRNA TSS isoforms of *MXI1*; the 5′ and 3′ UTR are colored white, and the position of uORFs is indicated by red arrows. **(d)** mRNA abundance of each *MXI1* mRNA TSS isoform estimated as transcript per million (TPM) from 5′ end-Seq data. Data presented are the average of the measurements of the three independent clones. **(e)** Similar to **b**, but the proportion of each *MXI1* mRNA TSS isoform in RCC4 cells is shown separately.

Importantly, some of the largest effects on translation were associated with alternate TSS usage (y-axis of Fig. 4a). Of these Max-interacting protein 1 (*MXI1*), an antagonist of Myc proto-oncogene^23^, manifest the most striking increase in translational efficiency upon *VHL* loss (Fig. 4a and b). 5′ end-Seq identified the three most abundant *MXI1* mRNA isoforms in RCC4 cells defined by alternative TSS usage (TSS1-3, Fig. 4c). TSS2 and TSS3 isoforms were the dominant isoforms in HIF repressed RCC4 VHL cells. However, the TSS1 transcript (which has been reported to be HIF1A dependent^24^) was strongly upregulated in *VHL*-defective RCC4 cells (Fig. 4d). Notably, TSS2 and 3 mRNA each contain an uORF that is excluded from TSS1 by alternative first exon usage (Fig. 4c). Consistent with the negative effects of uORFs on translation, polysome analysis indicated that the TSS1 mRNA isoform was much more efficiently translated than TSS2 and TSS3 isoforms (Fig. 4e). Thus, alternative TSS usage associated with *VHL* loss specifically upregulated the translationally more potent isoform, resulting in enhanced overall translation. Taken together, these findings indicate that whereas HIF has relatively modest overall effects on translation compared to its large effects on transcription, alternative TSS usage makes major contributions to altered translational efficiency amongst a subset of HIF-target genes.

### Differential sensitivity to mTOR inhibition amongst classes of HIF target gene

Since concurrent dysregulation of HIF and mTOR pathways is frequently observed in cancer and physiological stress, we sought to determine how HIF-dependent transcriptional regulation and mTOR dependent translational regulation interact. We first examined whether HIF pathway activation alters the translational response to mTOR inhibition. Comparison of changes in translational efficiency with mTOR inhibition in RCC4 VHL cells with those in RCC4 cells showed a close correlation, with the slope of the regression line being slightly less than 1 (Pearson’s *r* = 0.89, *p* < 10^-10^, slope = 0.85; Fig. 5a), indicating that mTOR inhibition regulates translation similarly regardless of HIF status. The effect of mTOR inhibition was slightly weaker in *VHL*-defective cells, probably reflecting a small negative effect of HIF1A on mTOR target mRNAs as outlined above.

**Fig. 5.**
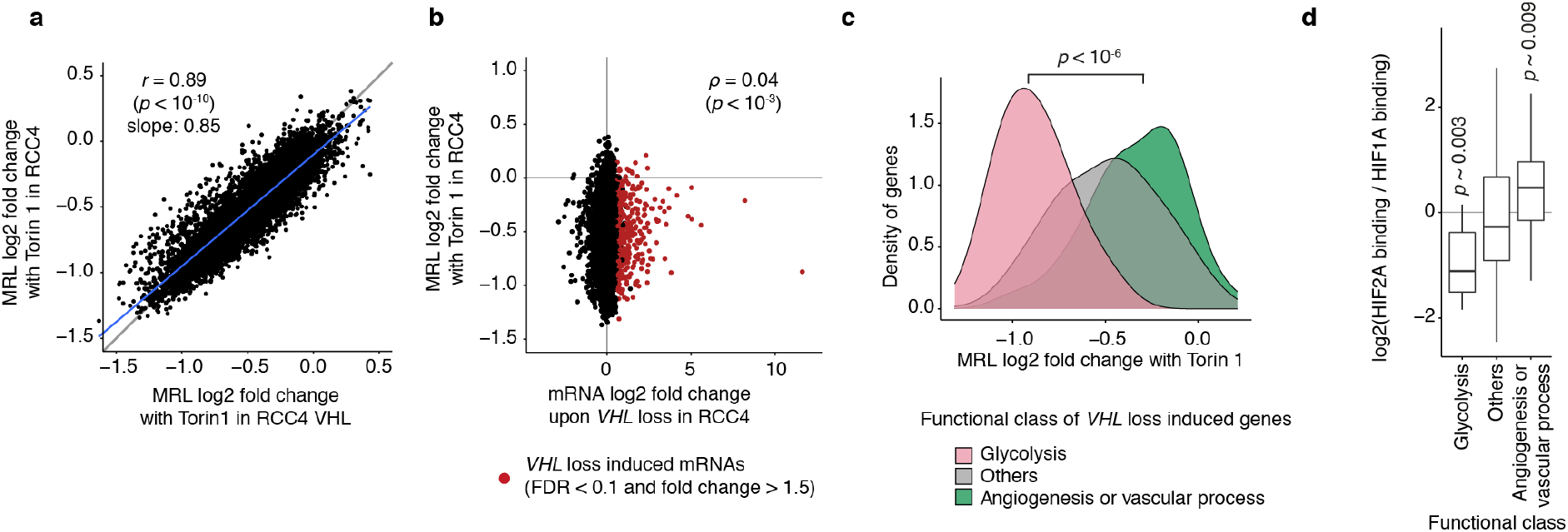
Differential sensitivity to translational inhibition by mTOR amongst HIF-regulated transcripts encoding proteins with different functions. **(a)** Comparison of effects of Torin 1 on translational efficiency of genes (expressed as log2 fold change in mean ribosomal load, MRL) in RCC4 VHL cells versus RCC4 cells. Pearson’s product moment correlation coefficient was used to assess the association (n = 8,429); the blue line indicates the linear model fit by ordinary least squares. **(b)** Comparison of the effect of Torin 1 on translational efficiency of genes with the effect of VHL on transcript abundance in RCC4 cells. Genes showing significant upregulation of mRNA abundance upon *VHL* loss are indicated in red (FDR < 0.1 and fold change > 1.5). Spearman’s rank-order correlation coefficient was used to assess the association (n = 8,580). **(c)** Analysis of changes in translational efficiency of genes produced by Torin 1 amongst the specified functional classes of genes whose mRNAs were induced by *VHL* loss. Functional classes were defined by gene ontology and KEGG orthology. The distributions are shown using kernel density estimation, and compared using the Mann-Whitney *U* test (n = 12 and 29 for glycolysis and angiogenesis or vascular process genes respectively). **(d)** Relative ratio of HIF2A and HIF1A binding at the nearest HIF binding sites to genes induced by *VHL* loss, amongst the specified functional class of genes. HIF2A and HIF1A binding across genome were analyzed by ChIP-Seq. The ratios within a functional class were compared against the ratios for all other genes using the Mann-Whitney *U* test (n = 12, 25, and 268 for glycolysis, angiogenesis or vascular process and others, respectively).

We then analyzed the relationship of HIF-dependent changes in transcription to mTOR-dependent changes in translation. Somewhat surprisingly we observed no overall association between the two regulatory modes (Spearman’s *ρ* = 0.04, *p* < 10^-3^; Fig. 5b). However, more detailed examination of the data revealed that distinct functional classes of mRNAs responded differently. Amongst transcripts that were induced in *VHL*-defective cells, those encoding glycolytic enzymes were hypersensitive to mTOR inhibition, whilst the translation of genes classified as involved in angiogenesis or vascular processes was much more resistant (*p* < 10^-6^, Mann-Whitney *U* test, Fig. 5c, Supplementary Fig. 8a and Supplementary Table 3). Indeed, the majority of glycolytic genes whose transcripts were upregulated in *VHL*-defective cells manifest significant translational repression by mTOR inhibition (Supplementary Fig. 8b). This indicates that full upregulation of glycolysis pathway requires both HIF and mTOR activity as would be predicted to occur in *VHL*-defective kidney cancer with mTOR hyperactivation^2^.

Of the two mTOR complexes it is widely accepted that mTORC1, regulates translation^9^. Interestingly, the protein level of HIF1A and HIF2A has been shown to be differentially regulated by mTORC1 and mTORC2. HIF1A is positively regulated by both mTORC1 and mTORC2 while HIF2A is dependent on only mTORC2 activity^25^, raising a question as to whether the HIF-induced, mTOR resistant genes that function in angiogenesis or vascular processes might be principally regulated by HIF2A and hence transcriptionally and translationally resistant to mTORC1 inhibition. To this end, we interrogated pan-genomic data on HIF binding^26^. In agreement with previous studies showing that genes encoding glycolytic enzymes are induced specifically by HIF1A^27^, HIF binding sites near this class of genes had a lower HIF2A/HIF1A binding ratio than other genes (*p* ∼ 0.003, Mann-Whitney *U* test, Fig. 5d). This contrasted with a higher HIF2A/HIF1A binding ratio for angiogenesis or vascular process genes induced in *VHL*-defective RCC4 cells (*p* ∼ 0.009, Mann-Whitney *U* test, Fig. 5d). Consistent with this, mRNAs of HIF-target angiogenesis or vascular processes genes were also upregulated to a greater extent than other HIF-target genes upon *VHL* loss in 786-O cells, which express only HIF2A (*p* ∼ 0.007, Mann-Whitney *U* test, Supplementary Fig. 8c). This suggests that they are primarily HIF2A targets, as well as being resistant to effects of mTOR inhibition on translation, consistent with a role for their increased protein expression in correcting a hypoxic and nutrient depleted environment.

## Discussion

Using a new technology to measure absolute translational efficiency of mRNAs resolved by their TSS, we have characterised the pan-genomic interplay of HIF and mTOR dependent transcriptional and translational regulation in *VHL*-defective kidney cancer cells. Importantly, the increased throughput of the technology and measurement of absolute translational efficiency enabled us to directly compare translational effects across the genome for a larger number of interventions than most studies to date. The direct measurement of absolute changes in translational efficiency is important since other genome-wide measurements such as ribosomal profiling generally refer separate measurements of mRNA sequencing reads and ribosome-protected sequencing reads to normalization values (total mapped reads) that are specific to each measurement. They therefore have to operate under an assumption of stability in the ratio of these normalization values, which unlikely to be valid in the face of a general action on translation such as with mTOR inhibition^13^.

Our analysis revealed that mTOR inhibition heterogeneously downregulates translation of a very wide variety of mRNAs and demonstrated the hypersensitivity of many genes encoding metabolic enzymes. This suggests a greater role for translational alterations in gene expression and metabolism in mTOR-dysregulated cancer than previously thought. The work also sheds new light on features of a transcript associated with hypersensitivity or resistance to mTOR inhibition.

Our findings confirmed that the HIF pathway primarily regulates transcription, but also revealed that HIF1A represses global translation moderately via mTOR and that HIF regulates the translation of a subset of genes bidirectionally through alternate TSS usage. Apart from these transcripts, we were surprised to find little or no evidence for HIF dependent up-regulation of translation in *VHL*-defective cells, given reports of a major role for HIF2A in promoting eIF4E2-dependent translation. The original studies demonstrated this action of HIF2A in hypoxia and in *VHL*-defective cells (786-O)^5, 6^ as were used in this study, but the effect size of HIF2A-dependent translational regulation was not compared with other interventions such as mTOR inhibition. Though we cannot exclude small effects on some targets, our findings indicate that, at least under the conditions of our experiments, the role of HIF2A/EIF4E2 in promoting translation is at best very limited.

Previous studies have reported that HIF inhibits mTOR activity through the transcriptional induction of antagonists of mTOR signalling^4, 28^, raising a question as to whether the use of mTOR inhibitors constitutes a rational approach to the treatment of *VHL*-defective cancer. Our comparative analysis of interventions revealed that the mTOR inhibition by HIF was very much weaker than that by pharmacological inhibition, offering a justification for this therapeutic approach.

To pursue this further, we compared transcriptional targets of HIF and translational targets of mTOR across the genome. Although little or no overall correlation was observed, these analyses revealed marked differences in mTOR sensitivity amongst HIF transcriptional targets, according to the functional classification of the encoded proteins. HIF1A-targeted genes encoding glycolytic enzymes were hypersensitive to mTOR whereas HIF2A-targeted genes encoding proteins involved in angiogenesis and vascular process were resistant to mTOR inhibition. Clinically approved mTOR inhibitors primarily target mTORC1^12^, and are therefore unlikely to affect HIF2A abundance^25^. Our results suggest that they are therefore unlikely to affect the expression of these classes of HIF2A-target gene. Recently a new class of drug that targets a pocket in the PAS domain of HIF2A to prevent dimerization with HIF1B subunits and hence blocks HIF transcriptional activity has shown promise in the therapy of *VHL*-defective kidney cancer^7, 8, 12^. Given that we observed few if any effects of HIF2A on translation, our results suggest that the combined use of these HIF2A transcriptional inhibitors, together with mTOR inhibitors, should therefore be considered as a rationale therapeutic strategy in this type of cancer.

## Methods

### Overview of the cell line and experimental conditions

*VHL*-defective kidney cancer cell lines, RCC4 and 786-O, cells were obtained from Cell Services at the Francis Crick Institute and were maintained in DMEM (high glucose, GlutaMAX Supplement, HEPES, Thermo Fisher Scientific, 32430100) with 1 mM sodium pyruvate (Thermo Fisher Scientific, 12539059) and 10% FBS at 37 °C in 5% CO_2_. Cells were confirmed to be of the correct identity by STR profiling and to be free from mycoplasma contamination.

Hypoxic incubation was performed using an InvivO_2_ workstation (Baker Ruskinn) in 1% O_2_ and 5% CO_2_ for 24h. To inhibit mTOR, cells were treated with 250 nM of Torin 1 (Cell Signaling Technology, #14379) for 2 hours.

An overview of the experimental interventions and the analyses is provided in Supplementary Table 1. Biological replicates are individual experiments using different clones derived from same cell line. All other replicates are defined as technical replicates.

### Genetic modification of cells

#### Lentiviral transduction

Reintroduction of *VHL* or the empty vector control was performed using lentiviral transduction. The expression vector (pRRL-hPGK promoter-VHL-IRES-BSD) containing the coding sequence and the last 6 nucleotides of the 5′ UTR of *VHL* (RefSeq ID, NM_000551) and the empty control vector (pRRL-SFFV promoter-MCS-IRES-BSD) were constructed from pRRL-SFFV promoter-MCS-IRES-GFP (provided by Professor Kamil R. Kranc, Queen Mary University of London). Lentiviruses were prepared from these plasmids, and RCC4 or 786-O cells were transduced with the viruses. Three or four clones of each of *VHL* or empty vector transduced RCC4 or 786-O cells were isolated using flow cytometry. These cells were maintained in DMEM (high glucose, GlutaMAX Supplement, HEPES) with 1 mM sodium pyruvate, 10% FBS and 5 µg/mL blasticidin (Thermo Fisher Scientific, A1113903) at 37 °C in 5% CO_2_. Throughout the study, empty vector transduced RCC4 or 786-O cells are referred as RCC4 or 786-O, and *VHL* transduced RCC4 or 786-O are referred as RCC4 VHL or 786-O VHL.

#### CRISPR/Cas9 mediated *HIF1B* or *EIF4E2* inactivation of 786-O cells

CRISPR/Cas9 mediated inactivation of *HIF1B* or *EIF4E2* was performed using the electroporation of gRNA-Cas9 ribonucleoprotein (RNP). The crRNAs with the following sequences were synthesized by Integrated DNA Technologies (Alt-R CRISPR-Cas9 crRNA): HIF1B, rGrArCrArUrCrArGrArUrGrUrArCrCrArUrCrArC EIF4E2 (g1), rGrUrUrUrGrArArArGrArUrGrArUrGrArCrArGrU EIF4E2 (g2), rGrGrUrCrCrCrCrArGrGrArCrGrUrArCrCrArUrG.

The *HIF1B* and *EIF4E2* (g2) gRNA sequences were designed by Integrated DNA Technologies (Hs.Cas9.ARNT.1.AD and Hs.Cas9.EIF4E2.1.AH respectively) whereas the EIF4E2 (g1) gRNA sequence was designed by an online tool developed by the Feng Zhang lab (https://crispr.mit.edu).

To prepare the gRNA, 100 µM of crRNA and 100 µM of tracrRNA (Integrated DNA Technologies, 14899756) were annealed in duplex buffer (Integrated DNA Technologies, 11-01-03-01) by incubating at 95 °C for 5 min then at room temperature for 30 min. Cas9-gRNA RNP was formed by mixing 10 µM of the annealed tracrRNA-crRNA and 16.5 µg of TrueCut Cas9 protein (Thermo Fisher Scientific, A36498) in PBS and incubating at room temperature for 30 min. The RNP was transfected into 786-O cells or 786-O VHL cells (pools of cells were used for HIF1B inactivation whereas clone 1 of each sub-line was used for EIF4E2 inactivation). Transfections were performed using a 4D-Nucleofector System (Lonza) with a SF Cell Line 4D-Nucleofector X Kit L (Lonza, V4XC-2024) and the EW-113 transfection programme, according to the manufacturer’s instructions. The transfected cells were cultured in DMEM (high glucose, GlutaMAX Supplement, HEPES) with 1 mM sodium pyruvate and 10% FBS at 37 °C in 5% CO_2_ for at least 3 days and single clones were isolated using flow cytometry. Inactivation of the target genes was confirmed both by Sanger sequencing of the gRNA target region using TIDE analysis^29^ and by immunoblotting.

### Immunoblotting

#### Protein extraction

Cells were grown on 6 cm dishes. Cells were washed with 3 mL of ice-cold PBS and then lysed by adding 150 µL of urea SDS lysis buffer (10 mM Tris-HCl pH 7.5, 6.7 M urea, 5 mM DTT, 10% glycerol, 1% SDS, 1x HALT protease and phosphatase inhibitor (Thermo Fisher Scientific, 78447), and 1/150x (v/v) of benzonase (Sigma-Aldrich, E1014-25KU)). The lysate was incubated at room temperature for 30 min before mixing with protein sample loading buffer (LI-COR Biosciences, 928-40004).

#### Immunoblotting

Proteins were separated by SDS-PAGE using a Mini-PROTEAN TGX Gel (4-15% or 8-16%, Bio-Rad Laboratories, 4561086 and 4561106, respectively) and transferred to Immobilon-FL PVDF Membrane (Sigma-Aldrich, IPFL00010). Proteins on the membrane were stained using a Revert 700 Total Protein Stain (LI-COR Biosciences, 926-11011) according to the manufacturer’s instructions. The data acquisition was performed using an Odyssey CLx system (LI-COR Biosciences) and the data were analysed using Image Studio software (LI-COR Biosciences). The membrane was blocked by incubating in TBS (20 mM Tris-HCl pH 7.6 and 137 mM NaCl) with 5% fat-free milk for 1 h shaking at room temperature. The membrane was incubated in (for anti-HIF2A antibody) Odyssey Blocking Buffer (PBS, LI-COR Biosciences, 927-40000) with 0.2% Tween 20 and 1/1,000 (v/v) primary antibody or (for other primary antibodies) TBST (TBS with 0.1% Tween 20) with 5% fat-free milk and 1/1,000 (v/v) primary antibody, shaking overnight at 4 °C. The membrane was washed three times with TBST and incubated in Odyssey Blocking Buffer (PBS for anti-HIF2A antibody and TBS for other primary antibodies, LI-COR Biosciences, 927-50000) with 0.2 % Tween-20, 0.01% SDS, and 1/15,000 (v/v) secondary antibody, shaking for 1 hr at room temperature. The membrane was washed three times with TBST and once with TBS.

#### Antibodies

The following antibodies were used for the western blotting analysis: (primary antibodies) anti-VHL (Santa Cruz Biotechnology, sc-135657), anti-HIF1A (BD Biosciences, 610959), anti-HIF2A (Cell Signaling Technology, #7096), anti-HIF1B (Cell Signaling Technology, #5537), anti-EIF4E2 (Proteintech, 12227-1-AP), anti-NDRG1 (Cell Signaling Technology, #9485), anti-SLC2A1 (Cell Signaling Technology, #12939), anti-EGFR (Santa Cruz Biotechnology, sc-373746), and anti-CA9 (Cell Signaling Technology, #5649); (secondary antibodies) anti-mouse IgG DyLight 800 (Cell Signaling Technology, #5257) anti-mouse IgG IRDye 680RD (LI-COR Biosciences, 925-68072), anti-Rabbit IgG IRDye 800CW (LI-COR Biosciences, 926-32213).

### Total RNA extraction

Cells were grown on 6 well plates or 6 cm dishes. Total RNA used for the analysis of unfractionated mRNAs was extracted from the cells using the RNeasy Plus Mini Kit (QIAGEN, 74136) according to the manufacturer’s instructions, except for technical replicate 2 of the samples from RCC4 cells (see Supplementary Table 1). For these samples, cells were lysed with 350 µL of Buffer RLT Plus (QIAGEN, 1053393), and total RNA was extracted from the lysate using an RNA clean and concentrator-25 kit (Zymo Research, R1018) with the following modification: 752.5 µL of preconditioned RNA binding buffer (367.5 µL of RNA binding buffer (supplied with an RNA clean and concentrator-25 kit), 367.5 µL of absolute EtOH and 17.5 µL of 20% SDS) was added to the cell lysate. After mixing, the material was loaded onto the column of an RNA clean and concentrator-25 kit, and the manufacturer’s instructions were followed for the remaining steps.

### HP5 protocol (polysome profiling)

#### Sucrose gradient preparation

Sucrose gradients were prepared in polyallomer tubes (Beckman Coulter, 326819) by layering 2.25 ml 50% sucrose in 1x polysome gradient buffer (10 mM HEPES pH 7.5, 110 mM potassium acetate, 20 mM magnesium acetate, 100 mM DTT, 40 U/mL RNasin plus (Promega, N2615), 20 U/mL SuperaseIn RNase Inhibitor (Thermo Fisher Scientific, AM2694) and 100 µg / mL cycloheximide (Sigma-Aldrich, C4859-1ML)) under 2.15 ml of 17% sucrose in 1x polysome gradient buffer. Each tube was sealed with parafilm, placed on its side, and kept in the horizontal position at 4 °C overnight to form the gradient^30^.

#### Cell lysis and fractionation

Cells were grown on 15 cm dishes. To arrest mRNA translation, the cells (∼80% confluency) were treated with 100 µg / mL cycloheximide for 3 min. The medium was removed, and the dish was placed on ice during the following steps. Cells were washed with 10 mL of ice-cold PBS with 100 µg / mL cycloheximide. Cells were then lysed by adding 800 µL of polysome lysis buffer (10 mM HEPES pH 7.5, 110 mM potassium acetate, 20 mM magnesium acetate, 100 mM potassium chloride, 10 mM magnesium chloride, 1% Triton X-100, 2 mM DTT, 40 U/mL RNase plus, 20 U/mL SuperaseIn RNase Inhibitor, 1x HALT Protease inhibitor (Thermo Fisher Scientific, 78438) and 100 µg/mL Cycloheximide). The cytoplasmic lysate was homogenised by passage through a 25G syringe needle 5 times. To remove debris, the lysate was centrifuged at 1,200*g* for 10 min at 4 °C and the supernatant was collected. This material was centrifuged again at 1,500*g* for 10 min at 4 °C and the supernatant was collected. The protein and RNA concentration were measured using 660nm Protein Assay Reagent (Thermo Fisher Scientific, 22660) with Ionic Detergent Compatibility Reagent (Thermo Fisher Scientific, 22663) and Qubit RNA BR Assay Kit (Thermo Fisher Scientific, Q10210) respectively.

Lysate was then normalized according to the protein concentration, and 500 µL of the normalized lysate overlaid on the sucrose gradient, as prepared above. The gradient was ultracentrifuged at 55,000 rpm for 55 min at 4 °C with max acceleration and slow deceleration using an Optima LE-80K Ultracentrifuge and SW55Ti rotor (Beckman Coulter). The sucrose gradient was fractionated according to the number of associated ribosomes as determined by the profile of the absorbance at 254 nm using a Density Gradient Fractionation System (Brandel, Model BR-188). The fractionated samples were then snap-frozen on dry ice.

#### External control RNA addition and RNA extraction

Equal amounts of external control RNA were added to the polysome fractionated samples after thawing the snap frozen samples on ice. Commercially available external control RNA, including the ERCC RNA Spike-In Mix-1 kit (Thermo Fisher Scientific, 4456740) that we used, does not have a canonical mRNA cap. This can influence the template switching reaction efficiency during the 5′ end-Seq library preparation (see below). Thus, the amount of external control RNA added to the polysome fractionated samples was determined by preliminary experiments, so as to result in a library containing around 0.1% of reads from the external control RNA.

RNA was extracted from 150 µL of the fractionated samples using an RNA clean and concentrator-5 kit (Zymo Research, R1016) using the same procedure to extract RNA from unfractionated cell lysate (described above), and eluted into 10 µL of water. For a subset of samples as indicated in Supplementary Table 1, half of the input volume was used, and RNA was eluted into 8 µL of water. The integrity of the purified RNA was confirmed using a Bioanalyzer (Agilent).

### 5′ end-Seq protocol (see Supplementary Fig 1a)

#### Primer sequences

The sequences of oligonucleotide primers used for 5′ end-Seq are summarised in Supplementary Table 4. All the primers were synthesized and HPLC purified by Integrated DNA Technologies.

The 5′ end-seq method involves the following steps.

#### Step 1: Reverse transcription and template switching

cDNAs with adapter sequences at both the 5′ and 3′ ends were generated from full length mRNAs using a combined reverse transcription and template switching reaction. The RT primers, containing an oligo (dT) sequence, were annealed to the poly A tail of mRNAs by incubating 4 µL of reaction mix (1.9 µL of extracted RNA, 1 µL of 10 mM dNTP, 0.1 µL of 20 U/µL SUPERaseIn RNase-Inhibitor and 1 µL of 10 µM RT primer) at 72 °C for 3 min and held at 25 °C. Then 1 µL of 10 µM template switching oligo (TSO), and 5 µL of RT reaction mix (2 µL of 5x RT buffer (supplied with Maxima H Minus Reverse Transcriptase), 2 µM of 5 M betaine, 0.25 µL of water, 0.25 µL of SUPERaseIn RNase-Inhibitor, and 0.5 µL of 200 U/µL Maxima H Minus Reverse Transcriptase (Thermo Fisher Scientific, EP0753)) were added to the reaction. The TSO contained an adapter sequence (the constant region annealed by the PCR primers), an index sequence (to identify the sample source of the cDNA), unique molecular identifiers (UMI), and three riboguanosines at the 3’ end (to facilitate template switching reaction^31^). The reverse transcription and template switching reactions were performed using a program of 25 °C for 45 min, 42 °C for 25 min, 47 °C for 10 min, 50 °C for 10 min, 85 °C for 5 min and held at 4 °C.

#### Step 2: Enzymatic degradation of primers and RNA

Preliminary experiments indicated that the degradation of unused primers using a single-stranded DNA specific 3′-5′ exonuclease, Exonuclease I, reduced primer dimer artefacts in the subsequent PCR amplification, whereas the degradation of RNA by RNase H improved the yield of cDNA library. Furthermore, it is important to degrade TSO because if unused TSO contaminates the cDNA library after multiplexing, it confounds the library indexing. Since we suspected that the TSO is resistant to Exonuclease I due to the riboguanosines at the 3’ end, the TSO contains three deoxyuridines (after the adapter sequence, index sequence and UMI) so that the TSO can be degraded by the combination of an enzyme cleaving DNA at a deoxyuridine and Exonuclease I. Importantly, this degrades all the TSO except the adapter which forms a high melting temperature duplex with the cDNA protecting the cDNA from Exonuclease I. All these reactions were performed in a single step by adding 2 µL of enzyme mix (1 µL of Thermolabile USER II (New England Biolabs, M5508L), 0.5 µL of Exonuclease I (New England Biolabs, M0293S), and 0.5 µL of RNase H (New England Biolabs, M0297S)) to the sample and incubating at 4°C for 1 s, 37 °C for 1 h, 80 °C for 20 min, and held at 4 °C.

#### Step 3: Limited-cycle PCR amplification

15 µL of PCR reaction mix (1.25 µL of 10 µM of each PCR primer 1 Fw/Rev, 12.5 µL of KAPA HiFi HotStart Uracil+ ReadyMix (Roche, KK2802) and 1.25 µL of water) was added to the RT reaction and limited-cycle PCR amplification was performed using a program of 98 °C for 3 min; 4 cycles of 98 °C for 20 s, 67 °C for 15 s and 72 °C for 6 min; 72 °C for 5 min; and held at 4 °C.

#### Step 4: Multiplexing and optimized PCR cycle amplification

After adding 37.5 µL of ProNex beads (Promega, NG2002) to each sample, up to 16 samples were multiplexed. The cDNA library was purified according to the manufacturer’s instructions, eluted into 42 µL of 10 mM Tris-HCl, pH 7.4, then re-purified using ProNex beads (1.5:1 v/v ratio of beads to sample) and eluted into 45 µL of 10 mM Tris-HCl, pH 7.4. The library was reamplified by preparing PCR reaction mix (20 µL of cDNA library, 25 µL of KAPA HiFi HotStart Uracil+ ReadyMix and 2.5 µL of 10 µM each PCR primer 1 Fw/Rev) and using a program of 98 °C for 3 min; 4-6 cycles (see below) of 98 °C for 20 s, 67 °C for 15 s and 72 °C for 6 min; 72 °C for 5 min; and held at 4 °C. The number of PCR cycle for each amplification was determined by a pilot experiment using quantitative PCR (qPCR) to ensure that the amplification was at the early linear phase. The amplified cDNA library was purified using ProNex beads as above, and eluted into 26 µL of 10 mM Tris-HCl, pH 7.4. The purified cDNA library was quantified using a Qubit dsDNA HS Assay Kit (Thermo Fisher Scientific, Q32851).

#### Step 5: tagmentation

Tagmentation with Tn5 transposase was performed on 90 ng aliquots of the cDNA library using an Illumina DNA Prep kit (Illumina, 20018704) according to the manufacturer’s instructions.

#### Step 6: PCR amplification of mRNA 5′ end library

The “tagmented” library was attached to the beads of an Illumina DNA Prep kit and limited-cycle PCR amplification was performed by adding 50 µL of the following reaction mix (2.5 µL of 10 µM each of the PCR primer 2 Fw/Rev, 20 µL of Enhanced PCR Mix (supplied with an Illumina DNA Prep kit), and 27.5 µL of water) and using a program of 68 °C for 3 min; 98 °C for 3 min; 3 cycles of 98 °C for 45s, 62 °C for 30s and 68 °C for 2 min; 68 °C for 1 min, and held at 10 °C. The PCR primers used here anneal to the TSO and an adapter added by tagmentation, and thus specifically amplify DNA fragments containing 5′ end of mRNAs. The amplified mRNA 5′ end library was purified using ProNex beads as above and eluted into 25 µL of 10 mM Tris-HCl (pH 7.4).

The mRNA 5′ end library was reamplified by preparing a PCR reaction mix (10 µL of the mRNA 5′ end library, 25 µL of KAPA HiFi HotStart ReadyMix, 2.5 µL of 10 µM each of PCR primer 3 Fw/Rev (containing i5 and i7 index sequences) and 12.5 µL of water) and using a program of 98 °C for 3 min; 5 cycles (cycle number determined by a pilot experiment to define the early linear phase, as described above) of 98 °C for 20s, 62 °C for 15s, and 72 °C for 30s; 72 °C for 5 min; and held at 4 °C. The mRNA 5′ end library was again purified using ProNex beads (1.4:1 v/v ratio of beads to sample) according to the manufacturer’s instructions, and eluted into 20 µL of 10 mM Tris-HCl, pH 7.4. The purified mRNA 5′ end libraries were multiplexed again and then sequenced on HiSeq 4000 (Illumina) using paired-end (2x 100 cycles) and dual-index mode.

### Overview of computational data analyses

The computational pipeline used for the data analysis is available from (https://github.com/YoichiroSugimoto/20210219_HP5_VHL_mTOR); the pipeline can generate most of plots and results using the sequence data as input; sequence data are available from ArrayExpress (HP5: E-MTAB-10689, 5′ end-Seq of total mRNAs: E-MTAB-10688). Data analyses were performed using R (4.0.0)^32^ and the following packages (data.table (1.12.8)^33^, dplyr (1.0.0)^34^, stringr (1.4.0)^35^, magrittr (1.5)^36^, and ggplot2 (3.3.1)^37^) were used throughout.

The following reference data were used to annotate the data; human genome: hg38, obtained via BSgenome.Hsapiens.UCSC.hg38^38^; human transcripts: RefSeq^39^ (GRCh38.p13) and GENCODE^40^ (GENCODE version 34: gencode.v34.annotation.gtf), these two reference data were combined and redundant GENCODE entries that have a corresponding RefSeq annotation were removed.

Prior to the high-throughput DNA sequencing data analysis, sequencing data from the technical replicates were concatenated. Data are presented as the mean value of the biological replicates.

TSS boundaries and their associated mRNA isoforms were identified by 5′ end-Seq of total (unfractionated) mRNAs. The TSSs assigned to a particular gene were those mapping within 50 base pairs of that gene locus, as specified by RefSeq and GENCODE. The abundance of the mRNA isoform associated with each TSS is the number of reads starting from that TSS. The gene-level mRNA abundance is the sum of these isoforms for the relevant gene.

### Statistics

The correlation of two variables was analysed with the cor.test function of R to calculate statistics based on Pearson’s product moment correlation coefficient or Spearman’s rank correlation coefficient. The difference of two distributions was tested using the two-sided Mann-Whitney *U* test (for two independent samples) or the two-sided Wilcoxon signed-rank test (for paired samples). To analyse the effect size of the Wilcoxon signed-rank test, the matched-pairs rank biserial correlation coefficient^41^ was calculated using the wilcoxonPairedRC function of the rcompanion package^42^.

Boxplots show the median (horizontal lines) and first to third quartile range (boxes) of data. Kernel density estimation was performed using the geom_density function of the ggplot2 package with the parameter, bw = “SJ”.

### Sequencing read alignment

#### Read pre-processing

The sequence at positions 1-22 of read 1 is derived from the TSO and was processed before mapping. First, the UMI located at positions 10-16 was extracted using UMI-tools^43^. Note that the UMI was not used in the analyses since we found that the diversity of UMI was not sufficient to uniquely mark non-duplicated reads. Next, the library was demultiplexed using an index sequence located at positions 1-8, after which the constant regions of the TSO located at position 9 and positions 17-22 were removed using Cutadapt^44^ with the parameters, -e 0.2 --discard-untrimmed.

#### Read alignment

The pre-processed reads were first mapped to cytoplasmic rRNAs (NR_023363.1 and NR_046235.1), mitochondrial ribosomal rRNAs (ENSG00000211459 and ENSG00000210082) and ERCC external control RNAs (https://www-s.nist.gov/srmors/certificates/documents/SRM2374_putative_T7_products_NoPolyA_v1.fasta) using Bowtie2 software^45^ with the following parameters, -N 1 --un-conc-gz. The unmapped reads were then aligned to the human genome (hg38: sequence obtained via BSgenome.Hsapiens.UCSC.hg38^38^) with the annotation described above using STAR software in two pass mode^46^ with the following parameters, --outFilterType BySJout -- outFilterMultimapNmax 1.

### Definition of TSS peaks and boundaries

To define clusters of TSS, we considered two widely used peak callers, paraclu^47^ and decomposition-based peak identification (dpi)^48^ software. Our preliminary analysis indicated that paraclu software was more accurate in determining total peak area whereas dpi was more accurate in resolving peaks within multimodal clusters. To obtain the most accurate resolution and quantification of TSS clusters, we therefore combined the strength of these programmes and included information from existing large-scale database using the following 4-step procedure.

#### (Step 1) Definition of cluster areas

Using the standard workflow of paraclu software on pooled data from normoxic cells, RCC4, RCC4 VHL, 786-O, and 786-O VHL, cluster areas of 5′ termini were identified.

#### (Step 2) Definition of TSS clusters within cluster areas

The cluster areas defined above were further resolved by combining above data with FANTOM5 data, and applying dpi software as was originally used for FANTOM5 to resolve *bona fide* sub-clusters within the current data. Internal sub-cluster boundaries were defined as the midpoint between adjacent dpi-identified peaks.

#### (Step 3) Quality controls and filters

Artefactual clusters of 5′ termini, potentially generated by internal TSO priming, were filtered on the basis of a low (<15%) proportion of reads bearing non-genomic G between the TSO and mRNA, as the template switching reaction commonly introduces such bases at the mRNA cap but not following internal priming^4^. Since mitochondrial mRNAs are not capped, these transcripts were filtered if not overlapping an annotated site.

A further filter was applied to remove TSS subclusters of low abundance mRNA isoforms whose biological significance is unclear; low abundance was defined as ≦10% of the most abundant mRNA isoform for the relevant gene in any of the analyses.

#### (Step 4) Final assignment of TSS boundaries

To provide the most accurate identification of the TSS peaks and their boundaries, the resolved and filtered peaks from Step 3 were mapped back onto the input cluster areas as defined in Step 1, boundaries being set at the midpoint between filtered peaks.

### Assignment of transcripts to TSS

To identify mRNA features that might affect translational efficiency, we used base specific information on 5′ termini and assembled paired-end reads starting from each TSS (StringTie software^49^) to define the primary structure of the 5′ portion of the transcript. We then used homology with this assembly to assign a full-length transcript from RefSeq and GENCODE. The CDS of the assigned transcript was then used for the analysis. In small number of cases, where this TSS was downstream of the start codon, we took the most upstream in-frame AUG sequence to redefine the CDS. The most abundant primary structure from each TSS and its CDS were then used in calculation of the association of mRNA features with mean ribosome load (see below). Details of this process are given at the computational pipeline.

### mRNA feature evaluation

Features within the mRNA (e.g. TOP motif, structure near cap) were evaluated at base specific resolution using the following formula:

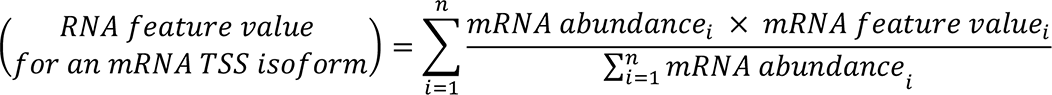

where *i* is a base position within the TSS, *n* is the linear sequence extent of the TSS, *mRNA feature value _i_* is the value of mRNA feature for the isoform transcribed from position *i*, and *mRNA abundance _i_* is the mRNA abundance of the isoform transcribed from position *i*. The values were rounded to the nearest integer; a rounded value of 0 being taken as the absence of the feature.

All non-overlapping uORFs, starting from an AUG, were identified using the ORFik package. Kozak consensus score was calculated by the kozakSequenceScore function of the ORFik package. Using the mode including G-quadruplex formation, the minimum free energy (MFE) of predicted RNA structures was estimated using RNALfold^50^. The MFE of RNA structures near the cap was that of the first 75 nucleotides. The MFE of the region distal to the cap was that of entire 5′ UTR minus the first 75 nucleotides. The position of a TOP motif was defined as the position of the 5′ most pyrimidine base, and its length was defined as that of the uninterrupted pyrimidine tract from that base.

### Functional annotation of genes

#### Functions

Functional classes of genes were defined by KEGG orthology^51^ as indicated by the following KEGG IDs; Transcription factors: 03000, Transcription machinery: 03021, Messenger RNA biogenesis: 03019, Spliceosome: 03041, Cytoplasmic and mitochondrial ribosome: 03011 (genes with the name starting with MRP and DAP3 were categorised as mitochondrial ribosomes), Translation factors: 03012, Chaperones and folding catalysts: 03110, Membrane trafficking: 04131, Ubiquitin system: 04121, and Proteasome: 03051; Glycolysis: hsa00010, Pentose phosphate pathway: hsa00030, TCA cycle: hsa00020, Fatty acid biosynthesis: hsa00061 and hsa00062, Fatty acid degradation: hsa00071, Oxphos: hsa00190, Nucleotide metabolism: hsa00230 and hsa00240, and Amino acid metabolism: hsa00250, hsa00330, hsa00220, hsa00270, hsa00260, hsa00340, hsa00310, hsa00360, hsa00400, hsa00380, hsa00350, hsa00290, and hsa00280.

Genes associated with angiogenesis or vascular process were defined by referencing to gene ontology (GO)^52^ database, GO:0003018: vascular process in circulatory system and GO:0001525: angiogenesis.

#### Analysis of existing literatures describing mTOR targets

To define known activities of mTOR via any mode of regulation (as indicated in Fig. 2c, first row) except translational regulation, we considered review articles by Saxton *et al.*^9^ and Morita *et al.*^53^. Known systematic translational down-regulation by mTOR inhibition (as indicated in Fig. 2c, second row) was defined from previous genome-wide studies performed with human cell lines^54, 55^. In Fig. 2c, a class of targets was defined as systematically downregulated if more than or equal to 10% of genes in the class were identified as mTOR hypersensitive in any of these previous studies^54, 55^.

### Analyses of differential mRNA expression upon *VHL* loss

The identification of differentially expressed genes and the calculation of log2 fold change in mRNA abundance upon *VHL* loss were performed using the DESeq2 package. Genes with an FDR < 0.1 and either log2 fold change > log2(1.5) or < −log2(1.5) were defined as upregulated or downregulated, respectively.

HIF-target genes (Supplementary Fig. 8c) were defined as those upregulated upon *VHL* loss in RCC4 cells. For this analysis, genes with very low expression in both 786-O and 786-O VHL cells, as identified by the DESeq2 package, were excluded from the analysis.

### Analysis of alternative TSS usage upon *VHL* loss

Genes manifesting alternative TSS usage upon *VHL* loss were identified using the approach described by Love *et al*^56^. Briefly, TSS for mRNA isoforms with very low abundance were first filtered out using the dmFilter function of the DRIMSeq package^57^ with the parameters, min_samps_feature_expr = 2, min_feature_expr = 5, min_samps_feature_prop = 2, min_feature_prop = 0.05, min_samps_gene_expr = 2, min_gene_expr = 20. The usage of a specific TSS relative to all TSS was then calculated by DRIMSeq with the parameter, add_uniform = TRUE.

The significance of changes in TSS usage upon VHL loss for a particular gene was analysed by the DEXSeq package. The FDR was calculated using the stageR package with a target overall FDR < 0.1. For genes with significant changes in VHL-dependent TSS usage, a *VHL*-dependent alternate TSS was selected as that showing the largest fold change upon *VHL* loss (FDR < 0.1) and a base TSS was selected as that showing the highest expression in the presence of VHL. In these calculations, the DESeq2 and apeglm package^58^ were used to incorporate data variance to provide a conservative estimate of fold change and standard error.

To provide the highest stringency definition, genes manifesting *VHL*-dependent alternative TSS usage were further filtered by the proportional change > 5%, the absolute fold change > 1.5, and the significance of the difference in fold change between the alternate TSS and the base TSS (assessed by non-overlapping 95% confidence intervals).

For the comparative analysis of the *VHL*-dependent alternate TSS usage in various conditions (Supplementary Fig. 6), genes with very low expression that did not meet a criterion of 20 read counts in more than 1 samples were excluded.

### Calculation of mean ribosome load

Mean ribosome load was calculated using the following formula:

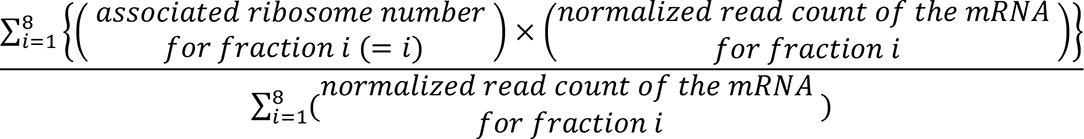

The mRNA abundance values for each polysome fraction were normalized by the read count of the external control using the estimateSizeFactors faction of the DESeq2 package. Very low abundance mRNAs that did not meet a criterion of 6 read counts in more than 6 samples were excluded.

### Statistical analysis of differences in polysome distribution

#### VHL-dependent alternate TSS mRNA isoforms

The significance of changes in the polysome profile of VHL-dependent alternate mRNA isoforms was determined by reference to all other isoforms from same gene by considering the ratio of mRNA abundances as a function of polysome fraction using the DEXSeq package^59^. The false discovery rate (FDR) was calculated using the stageR package^60^ with the target overall FDR < 0.1.

#### Differentially translated mRNA isoforms from same gene

In analysis of two most differentially translated mRNA isoforms transcribed from same gene (for Fig. 1f), each of these isoforms was censored for statistically significant differences from all other isoforms of same gene using the same analysis as above.

#### Changes in response to mTOR inhibition

To identify genes that were hypersensitive or resistant to mTOR inhibition, genes manifesting a significant change in polysome distribution upon mTOR inhibition, compared to the population average, were first identified using the DESeq2 package^59^ with the internal library size normalization and the likelihood ratio test. The genes with a significant change (FDR < 0.1) were classified as hypersensitive or resistant to mTOR inhibition if the log2 fold change of the mean ribosome load was lower or higher than the median of all expressed genes.

### Simulation of changes in translational efficiency with omitting a parameter

Mean ribosome load log2 fold change of a gene upon *VHL* loss can be expressed by the following formula:

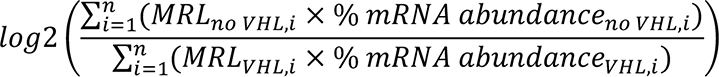

In this formula, *i* is mRNA isoform i (out of n mRNA isoforms), *MRL _no VHL or VHL, i_* is the mean ribosome load of isoform i in RCC4 or RCC4 VHL cells, and *% mRNA abundance _no VHL or VHL, i_* is the percentage abundance of isoform i relative to that of all isoforms in RCC4 or RCC4 VHL cells.

To assess the contribution of alternative TSS usage to changes in mean ribosome load of a gene, we tested a simulation which omitted the VHL-dependent changes in translational efficiency within each mRNA isoform using the following formula:

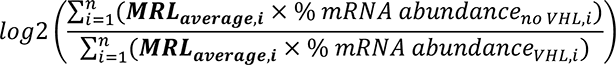

In this formula, *MRL _average, i_* is the combined average of *MRL _no VHL, i_* and *MRL _VHL, i_* as defined above. When values for either of *MRL _no VHL, i_* and *MRL _VHL, i_* are missing, these values are excluded from the calculation of the average.

To assess the contribution of VHL-dependent changes in translational efficiency within each mRNA isoform to changes in mean ribosome load of a gene, we tested a simulation which omitted the VHL-dependent changes in TSS usage using the following formula:

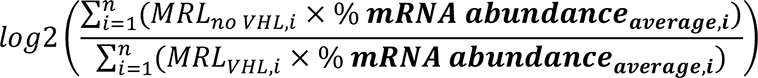

In this formula, *% mRNA abundance _average, i_* is the combined average of *% mRNA abundance_no VHL, i_* and *% mRNA abundance _VHL, i_* defined above. When values for either of *MRL _no VHL, I_* and *MRL _VHL, i_* are missing, these genes were excluded from the analysis.

### Generalized additive model to predict mean ribosome load

A generalized additive model was used to predict mean ribosome load of mRNAs from the preselected mRNA features. To test the model, a cross validation approach was deployed to predict the MRL of the top 50% expressed genes on 4 randomly selected chromosomes, which were excluded from the training data used to derive the model. To provide an accurate estimate of the model’s performance, this process was repeated 10 times and the median value of the coefficient of determination (*R^2^*) was calculated.

For model construction, the gam function of the mgcv package^61^ of R was used deploying thin plate regression splines with an additional shrinkage term (with the parameter, bs = “ts”) and restricted maximum likelihood for the selection of smoothness (with the parameter, method = “REML”). The analysis was restricted to mRNAs with 5′ UTR length longer than 0 nts and CDS length longer than 100 nts; 5′ UTR and CDS length were log10 transformed and MFE of RNA structures were normalized by the segment length [nucleotides].

### Principal component analysis

Library size normalization and a variance stabilizing transformation were applied to the mRNA abundance data using the vst function of the DESeq2 package^62^ with the parameter, blind = TRUE. Principal component analysis of the transformed data was performed for genes showing the most variance (top 25%) using the plotPCA function of the DESeq2 package.

### GO or KEGG orthology enrichment analysis

GO or KEGG orthology enrichment analysis of the selected set of genes compared to all the expressed genes in the data was performed using g:Profiler^63^ through the gost function of the R interface, the gprofiler2 package.

### Analysis of HIF2A/HIF1A binding ratio near VHL-regulated genes

HIF1A and HIF2A ChIP-Seq data from Smythies *et al.*^26^ were used to analyse HIF binding sites across the genome. HIF1A or HIF2A binding sites were defined as the overlap of the peaks identified by ENCODE ChIP-Seq pipeline (https://github.com/ENCODE-DCC/chip-seq-pipeline2) and those by MACS2 software^64^. For this purpose, the ChIP-Seq reads were aligned to the human genome using Bowtie2 software and the aligned reads were analysed by ENCODE ChIP-Seq pipeline to identify the peaks. The blacklist filtered and pooled replicate data generated by the pipeline were analysed by MACS2 software with the following parameters (callpeak -q 0.1 --call-summits). The position of the binding sites was defined as the position of the hypoxia response element (HRE, RCGTG sequence) closest to the peak summits identified by MACS2 software. If the binding site did not contain an HRE within 50 bp of the peak summit, it was filtered out. Data on HIF1A and HIF2A binding as defined above were merged, and HIF2A/HIF1A binding ratio was estimated using the DiffBind package^65^ with the parameters, minMembers = 2 and bFullLibrarySize = FALSE.

For the calculation of the distances between HIF binding sites and VHL-regulated genes (censored by 500 kb^26^), the position of the HIF bindings site was defined by that of the most abundant isoform at the site, and the position of the gene was defined by that of the TSS manifesting the greatest regulation by VHL.

## Acknowledgement

We thank the advanced sequencing facility team at the Francis Crick Institute for Illumina HiSeq sequencing; Ratcliffe group members for support and discussion; and Matthew Cockman for critical reading of the manuscript. This work was supported by the Francis Crick Institute which receives its core funding from Cancer Research UK (FC001501), the UK Medical Research Council (FC001501), and the Wellcome Trust (FC001501). PJR is also supported as a distinguished scholar of the Ludwig Institute for Cancer Research and by the Wellcome Trust (106241/Z/14/Z). For the purpose of Open Access, the author has applied a CC BY public copyright licence to any Author Accepted Manuscript version arising from this submission.

## Author contribution

Y.S. and P.J.R. conceived the project. Y.S. performed experiments and data analysis. Y.S. and P.J.R. contributed to the interpretation of the data. Y.S. and P.J.R. wrote the manuscript.

## Supplementary Tables

**Supplementary Table 1** Overview of high-throughput DNA sequencing based experiments

**Supplementary Table 2** VHL dependent alternative TSS genes and the translational effect

**Supplementary Table 3** mTOR sensitivity of HIF target genes regulating glycolysis, angiogenesis, or vascular process

**Supplementary Table 4** Primer sequences

## Supplementary Notes

Summary of exact n and *p* values for the analyses displayed in figures.

---

**Supplementary Fig. 1.**
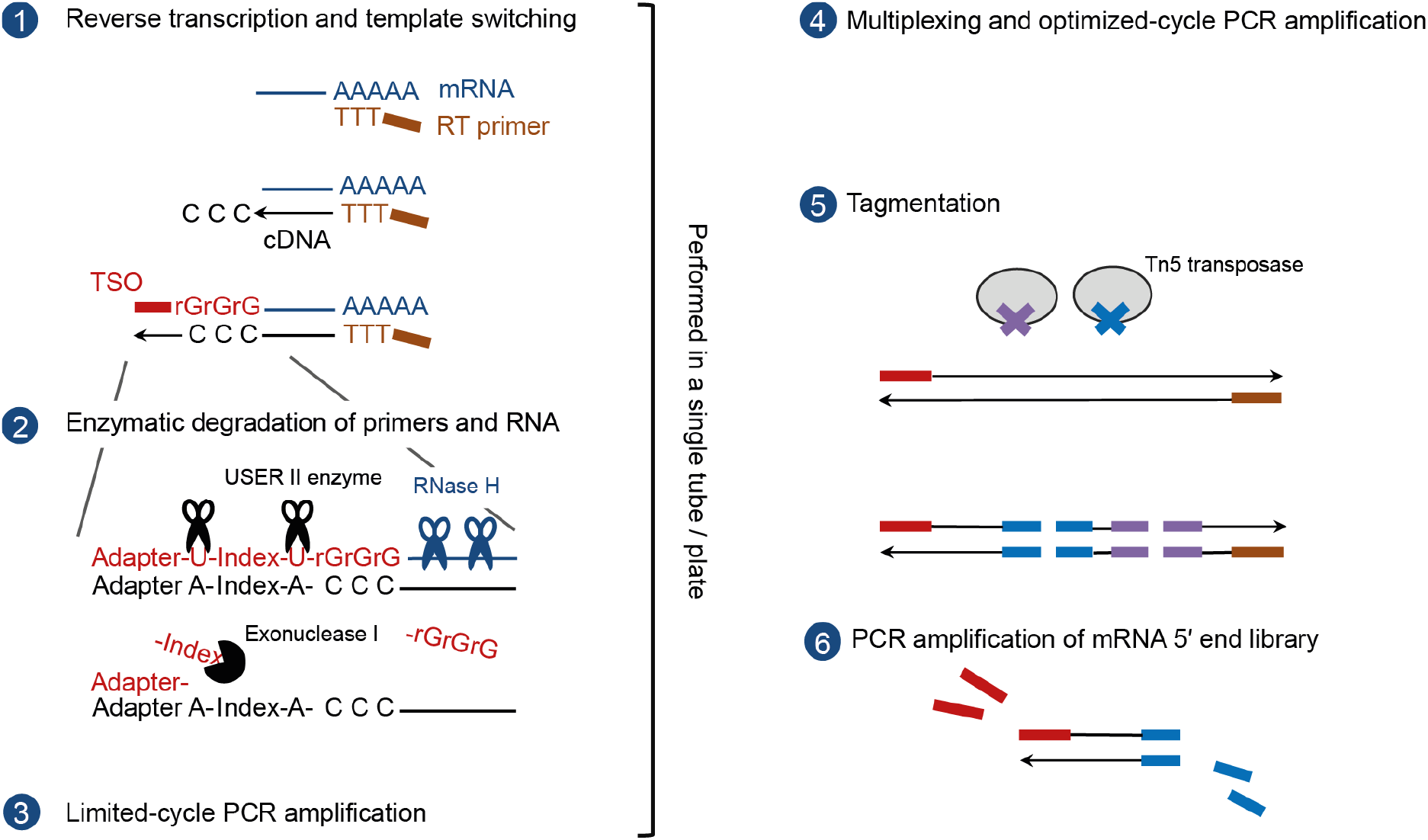
Overview of 5′ end-Seq protocol. Schematic representation of the 5′ end-Seq protocol (see also Methods). 1. The reverse transcription is primed with an adapter containing an oligo (dT) sequence. The reverse transcriptase used for 5′ end-Seq adds additional non-templated cytidine residues beyond the cap, to the 3′ end of the cDNAs. This polycytidine sequence anneals to a polyriboguanosine sequence contained in the template switch oligo (TSO), and the reverse transcriptase switches the template from the mRNA to the TSO to add the complementary sequence of the TSO at the 3′ end of the cDNAs. An indexing sequence contained in the TSO to identify the sample source of the cDNAs is reverse transcribed in this process. 2. Unused primers and RNA are degraded using the combination of a single-stranded DNA specific exonuclease (Exonuclease I), an enzyme cleaving DNA at deoxyuridine (Thermolabile USER II enzyme), and RNase H. This step leaves the adapter sequence of the TSO (the constant region) annealing to the cDNA due to the high melting temperature of this duplex, which protects the cDNA from Exonuclease I. 3. The full-length cDNA library is amplified using limited cycle PCR amplification. 4. The libraries from different samples are multiplexed and the multiplexed libraries are amplified using PCR and an optimized cycle number. 5. Amplified libraries are fragmented and adapter tagged using tagmentation. 6. mRNA 5′ end library suitable for high-throughput DNA sequencing is generated using PCR amplification with primers annealing to the TSO and the appropriate tagmentation adapter.

**Supplementary Fig. 2.**
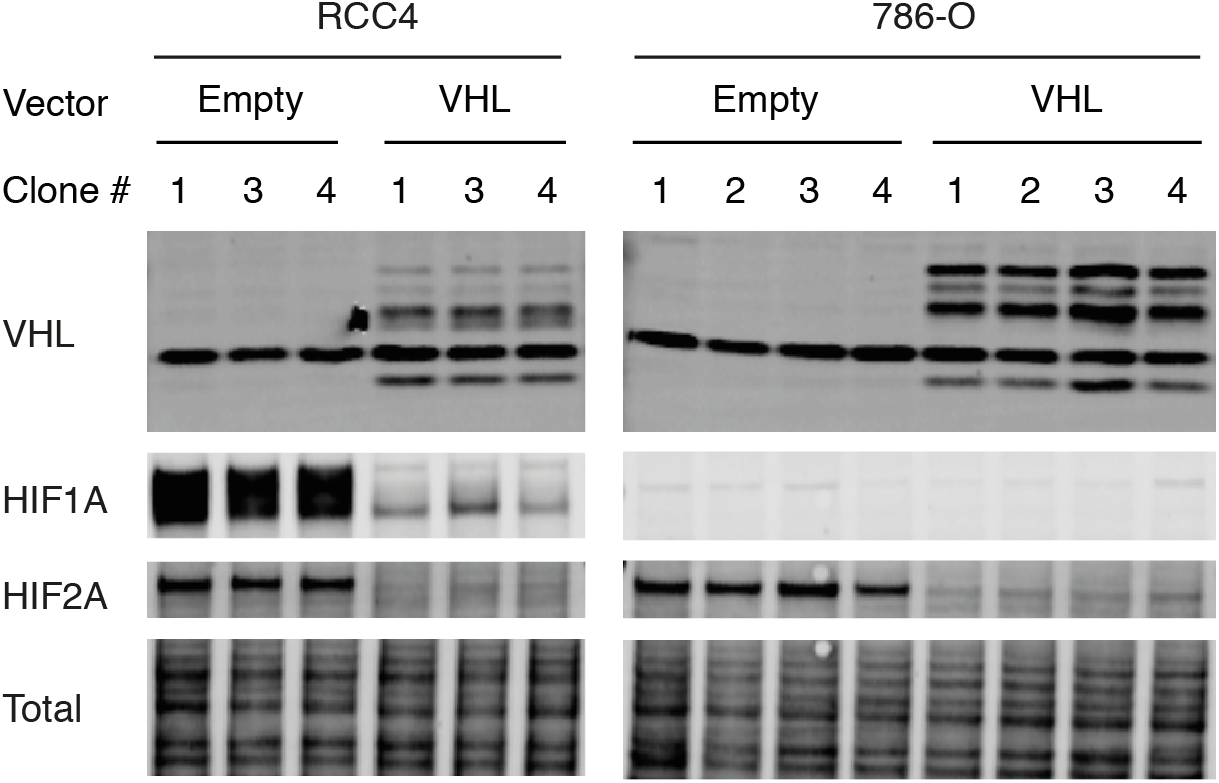
Establishment of cell lines. Immunoblotting analysis of RCC4 or 786-O cells re-expressing either wild type VHL or empty vector alone. The successful reintroduction of VHL was confirmed by the expression of VHL protein and degradation of HIF1A and/or HIF2A protein. Similar protein loading across lanes was confirmed by total protein staining.

**Supplementary Fig. 3.**
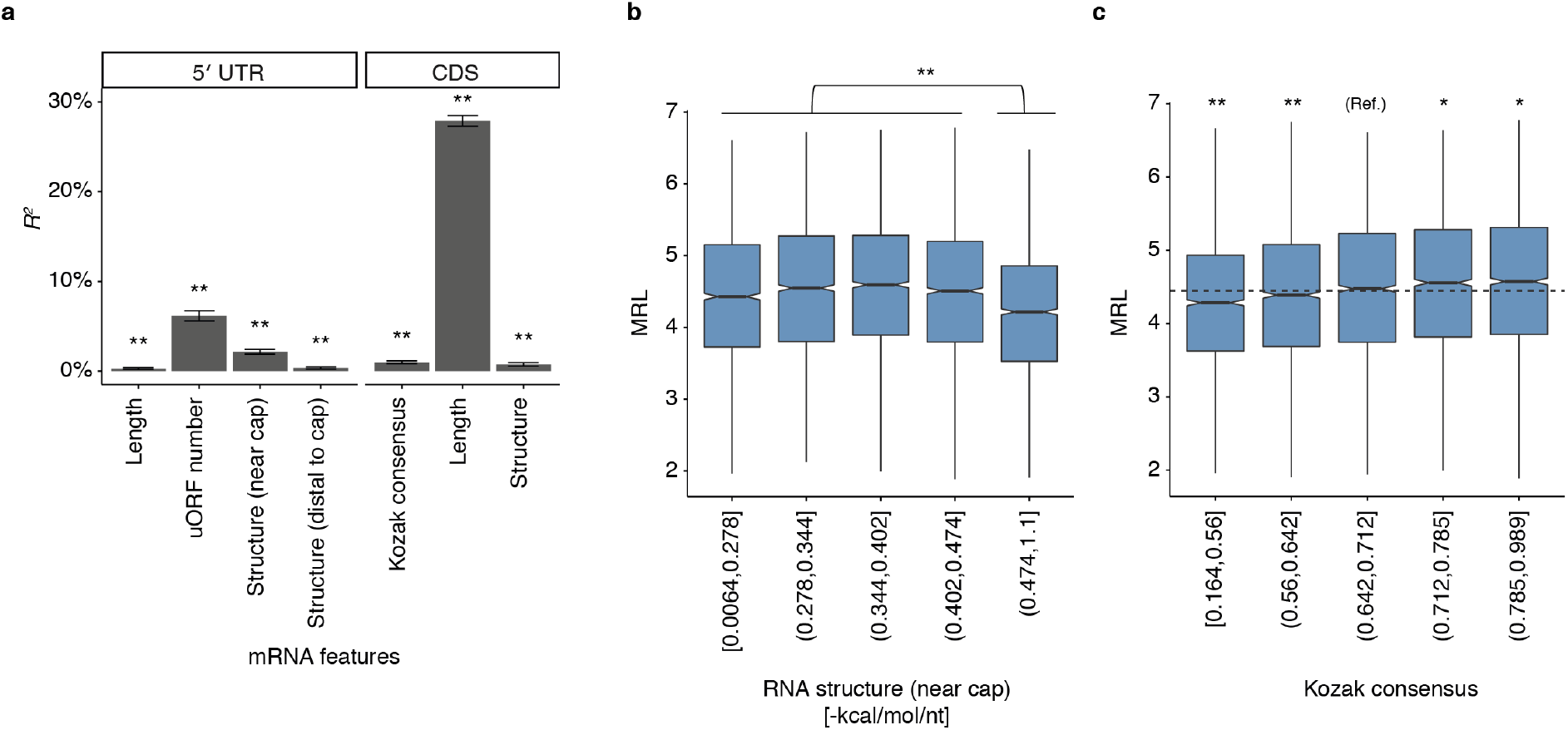
mRNA features predicting mean ribosome load. **(a)** Proportion of variance in mean ribosome load (MRL) between mRNAs that is explained by a single mRNA feature (expressed as *R^2^*) using a generalized additive model (mean±s.e.m of 10 iterations of cross validation). The significance of mRNA features in predicting MRL was determined by the Wald test. Length, log10 sequence length (nucleotides, nts); Structure (near cap, first 75 nts; distal to cap, rest of the 5′ UTR), inverse of minimum folding energy per nucleotide of predicted RNA structures; Kozak consensus, match score to the consensus Kozak sequence. The analysis identified that CDS length, uORF number, stability of RNA structures near cap, and Kozak consensus score were the four most predictive features. **(b)** MRL as a function of the stability of RNA structures near cap. mRNAs were ranked by their RNA structural stability, and split into 5 groups according to the rank (i.e. the rightmost group contains the top 20% mRNAs with the most stable RNA structures); the intervals of the stability are indicated on the x-axis. The distribution of MRL for mRNAs with less stable structures was compared with the most stable group (0.471 to 1.1 -kcal/mol/nt) using the Mann-Whitney *U* test. **(c)** Similar to **b**, but MRL as a function of Kozak consensus score. The median value of MRL for all mRNAs is shown by a dashed line. The distribution of MRL for mRNAs with the indicated Kozak consensus score was compared to that with the score of 0.642 to 0.712, using the Mann-Whitney *U* test. *: *p* < 0.05, ** *p* < 0.005. *P* values were adjusted for multiple comparisons using Holm’s method. Details of the sample sizes and exact *p* values are summarized in Supplementary Notes.

**Supplementary Fig. 4.**
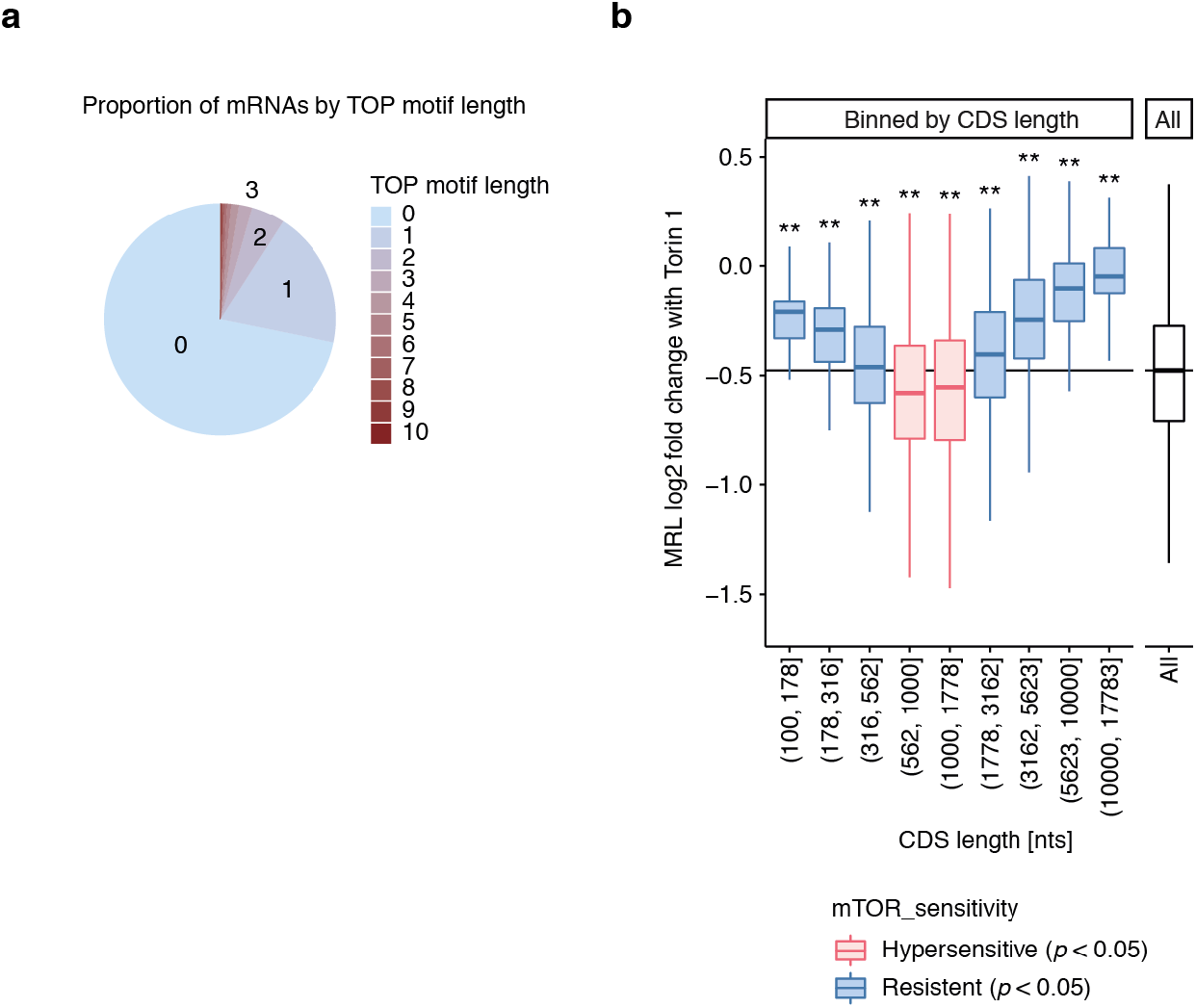
HP5 refined mRNA features influencing the mTOR sensitivity of mRNAs. **(a)** Proportion of mRNAs as a function of the TOP motif length (n = 9,589). **(b)** Box plots showing changes in translational efficiency of mRNAs with Torin 1 (log2 fold change mean ribosome load, MRL) as a function of CDS length. Responses of mRNAs with the indicated CDS length were compared against responses of all other mRNAs using the Mann-Whitney *U* test; classes more downregulated or less downregulated compared to all other mRNAs (i.e. hypersensitive or resistant to mTOR inhibition, *p* < 0.05) are colored red or blue respectively. *: *p* < 0.05, ** *p* < 0.005. *P* values were adjusted for multiple comparisons using Holm’s method. Details of the sample sizes and exact *p* values for (b) are summarized in Supplementary Notes.

**Supplementary Fig. 5.**
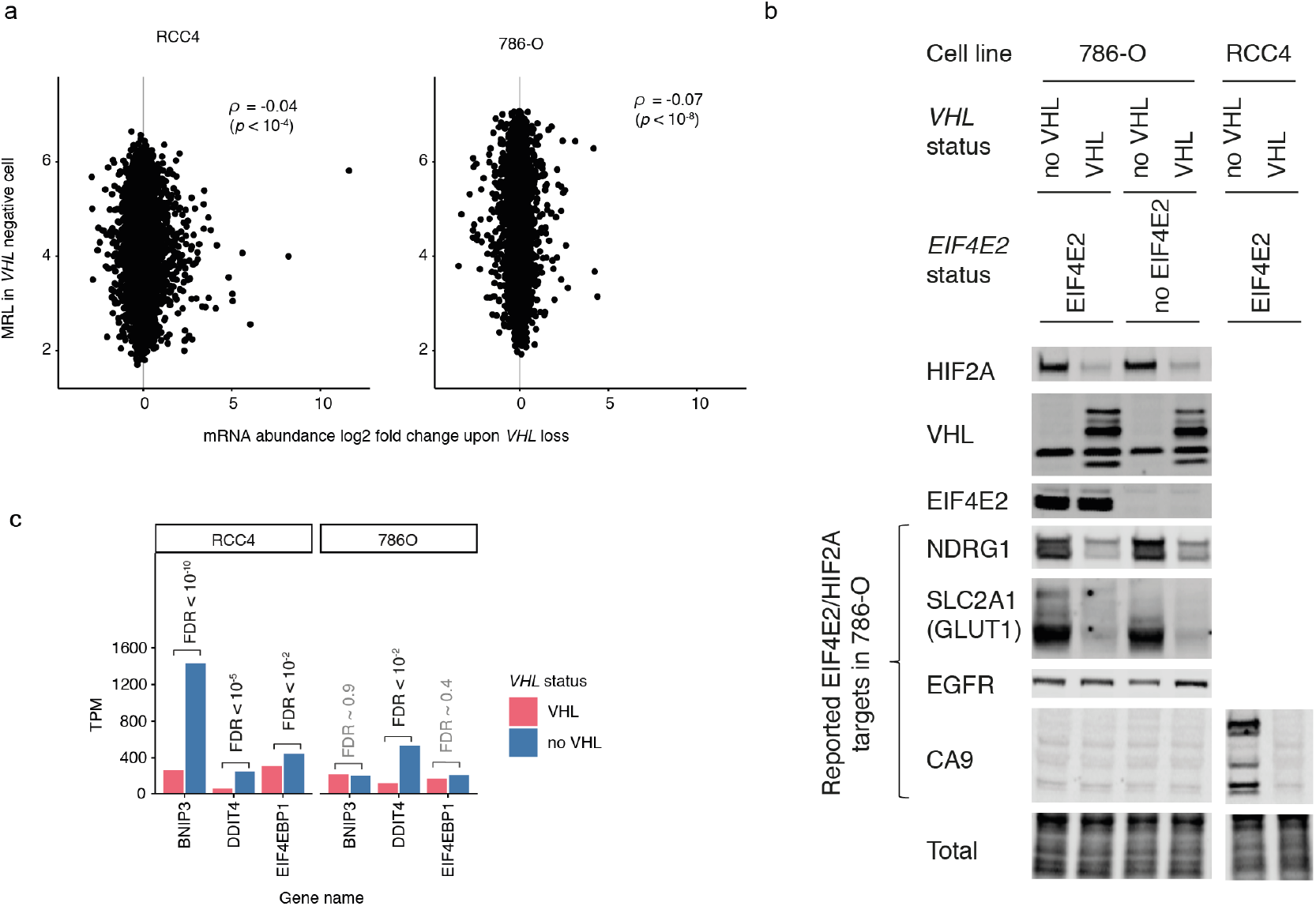
Effects of VHL on the efficiency of translation. **(a)** Scatter plots comparing changes in mRNA abundance of genes upon *VHL* loss with the mean ribosome load (MRL) in RCC4 and 786-O cells. Spearman’s rank-order correlation was used to assess the association (n = 9,493 and 8,065 for RCC4 and 786-O respectively). The absence of correlation indicates that genes induced by *VHL* loss were not preferentially translated upon induction compared to genes that are not induced by VHL. **(b)** Immunoblotting analysis of four previously reported EIF4E2/HIF2A target genes in 786-O cells^5, 6^ as a function of VHL and EIF4E2. HIF2A induction did not alter EGFR protein abundance. *EIF4E2* inactivation did not alter NDRG1 and SLC2A1 protein abundance in the presence of HIF2A indicating that the contribution of EIF4E2/HIF2A pathway to the regulation of the translational efficiency of these genes is at most very limited. Although CA9 was reported to be an EIF4E2/HIF2A target in 786-O cells^6^, CA9 protein expression could not be detected in 786-O cells in agreement with previous studies showing that CA9 is transcriptionally induced by HIF1A but not by HIF2A^66^. **(c)** Analysis of changes in mRNA abundance of negative regulators of mTOR pathway upon *VHL* loss. mRNA abundance was measured as transcripts per million (TPM) and the significance of differential expression was assessed using the Wald test (n = 3 and 4 for each condition in RCC4 and 786-O respectively).

**Supplementary Fig. 6.**
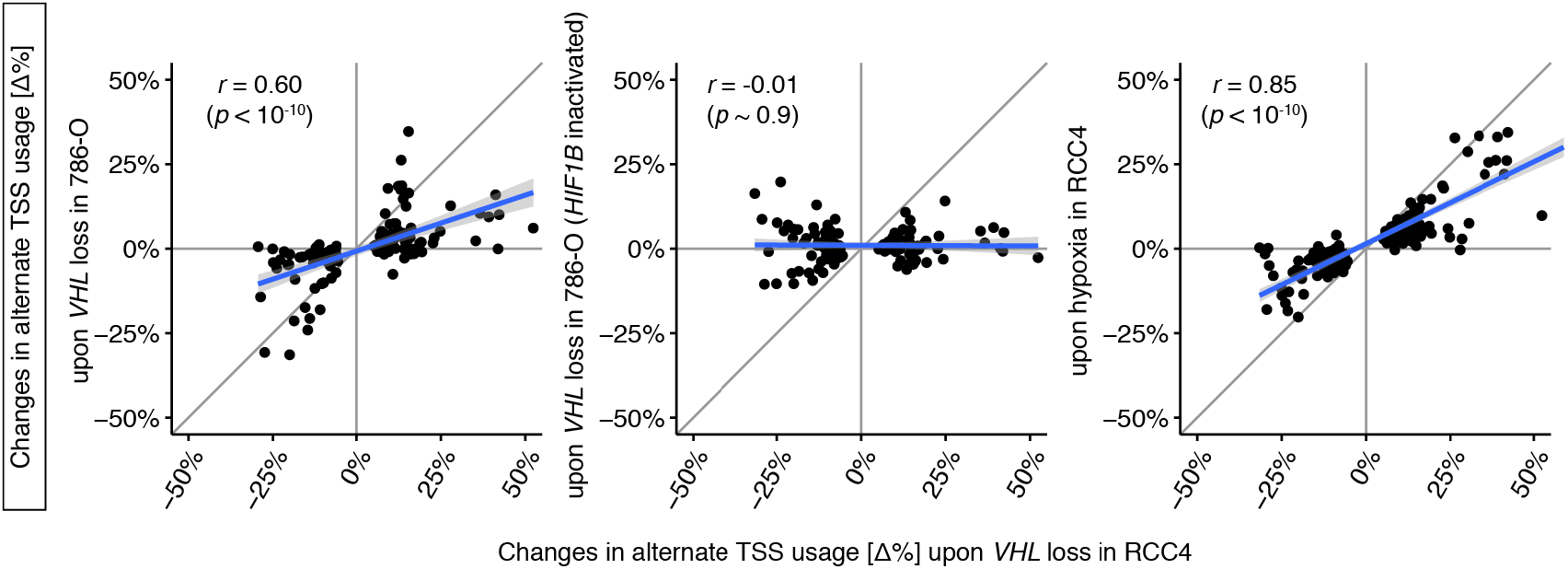
Alternate TSS usage in *VHL*-defective and hypoxic cells. Correlations between VHL-dependent changes in alternative TSS usage in RCC4 (x-axis) and such changes in other cells or conditions (y-axes); panels show correlations with *VHL* loss in 786-O cells (left); with *VHL* loss in *HIF1B* inactivated 786-O cells (middle) and with hypoxia (1% O_2_ for 24 h) in RCC4 VHL cells (right). Genes with too little mRNA expression for quantitative analysis were excluded from the analyses (see Methods). Pearson’s product moment correlation coefficient was used to assess the associations (n = 124, 126, and 148 for the respective comparisons).

**Supplementary Fig. 7.**
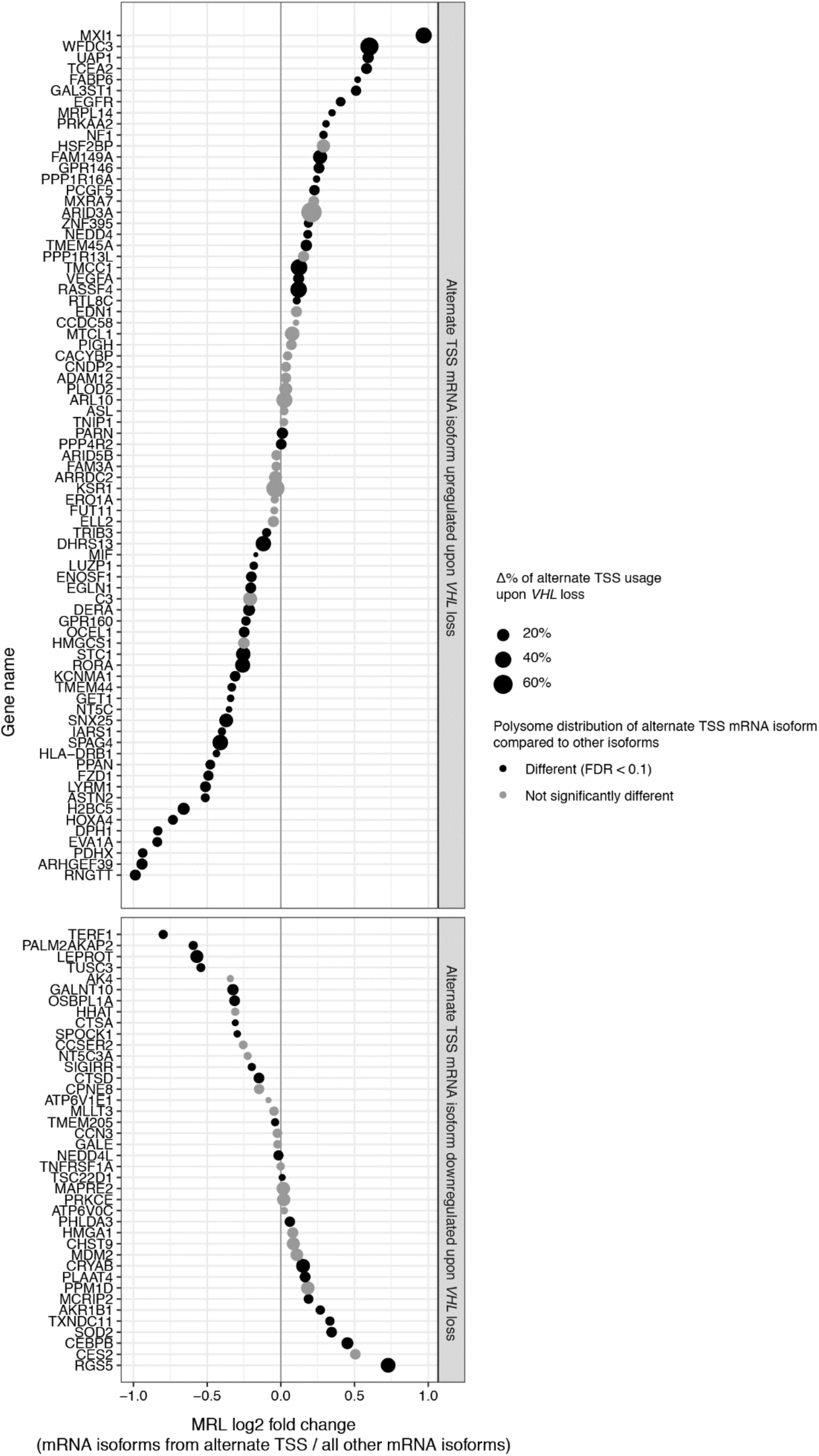
VHL dependent alternate TSS usage generates mRNAs with an altered translational efficiency. The plot shows the differences in translational efficiency (expressed as mean ribosome load, MRL) between mRNA isoforms that are generated from VHL-dependent alternative TSSs and all other isoforms transcribed from the same gene. Data are for RCC4 cells. Genes are sorted by log2 fold difference in MRL; significant differences in polysome distribution (FDR < 0.1) are indicated by black colouring. The magnitude of changes in alternative TSS usage is shown by the size of point. Genes with too little alternate TSS isoform expression for MRL calculation were excluded from the analysis (see Methods).

**Supplementary Fig. 8.**
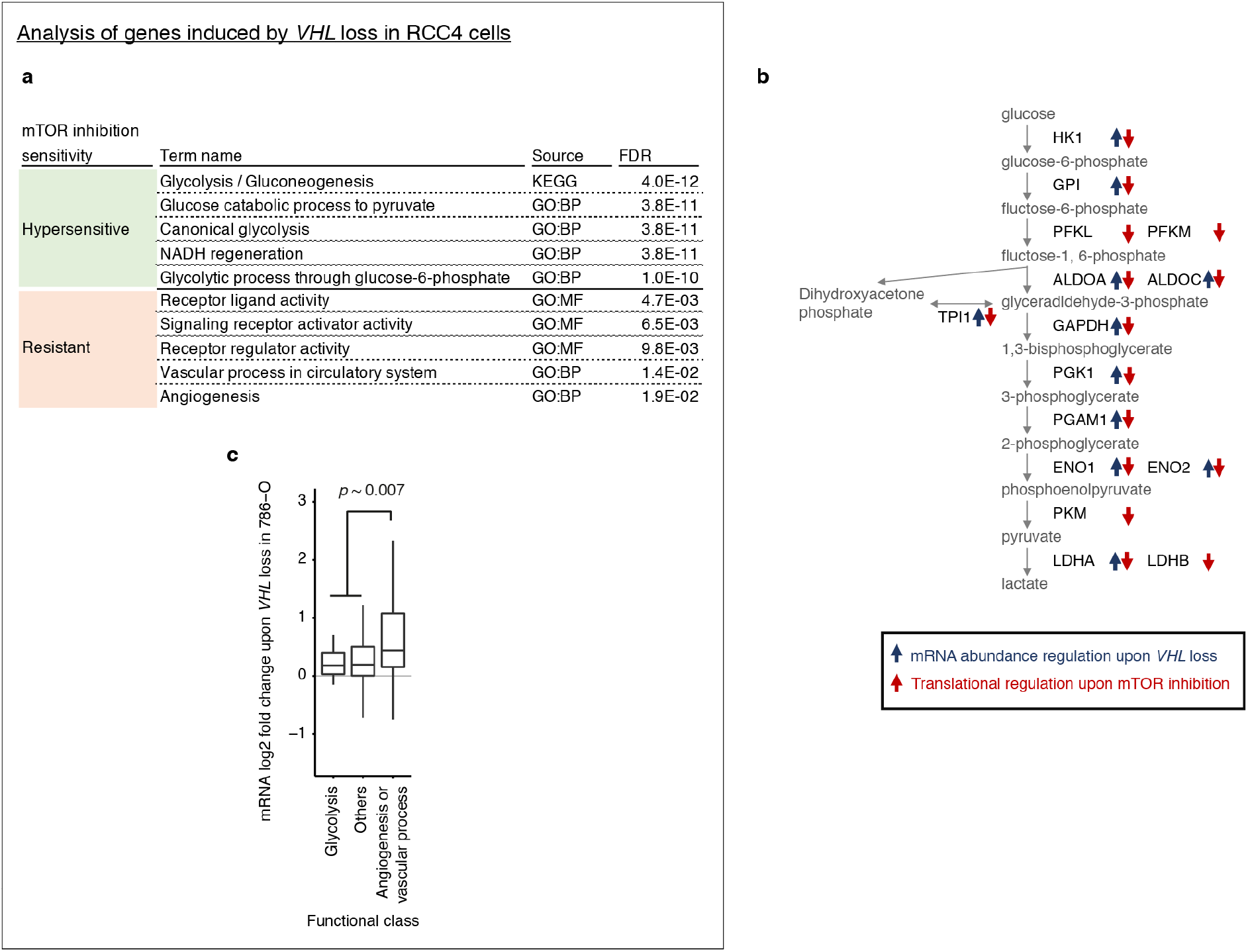
Differential sensitivity to mTOR inhibition amongst functional groups of transcripts induced upon *VHL* loss. **(a)** Gene set enrichment analysis among genes induced upon *VHL* loss (FDR < 0.1 and mRNA fold change > 1.5) and either hypersensitive (green) or resistant (pink) to mTOR inhibition in RCC4 cells (see Methods for definition). The top 5 the most enriched gene ontology or KEGG orthology terms are shown. n = 124 and 122 for mTOR hypersensitive and resistant genes were considered for the analysis. (b) Schematic showing inverse relationship of changes in mRNA abundance upon *VHL* loss (as defined above) and changes in translational efficiency upon mTOR inhibition (see Methods for definition) for genes encoding glycolytic enzymes in RCC4 cells. (c) Boxplots showing changes in mRNA abundance of HIF-target genes upon *VHL* loss in 786-O cells (i.e. induction of HIF2A) as a function of the encoded protein function. The changes in mRNA abundance of angiogenesis or vascular process genes were compared against those of all other HIF-target genes using the Mann-Whitney *U* test (n = 27 and 305 respectively).

