## Supplementary Notes for "Isoform resolved measurements of absolute translational efficiency define interplay of HIF and mTOR dysregulation in kidney cancer"

Summary of exact n and *p* values for the analyses displayed in figures.

**Fig. 1**

(d)

| CDS length | n | *p* |
|---|---|---|
| (100, 178] | 41 | < 10E-10 |
| (178, 316] | 468 | < 10E-10 |
| (316, 562] | 1,344 | < 10E-10 |
| (562, 1000] | 3,082 | < 10E-3 |
| (1000, 1778] | 4,100 | Reference |
| (1778, 3162] | 2,309 | < 10E-10 |
| (3162, 5623] | 731 | < 10E-10 |
| (5623, 10000] | 162 | < 10E-10 |
| (10000, 17783] | 30 | < 10E-5 |

(e)

| uORF number | n | *p* |
|---|---|---|
| 0 | 6,794 | Reference |
| 1 | 2,706 | < 10E-10 |
| 2 | 1,325 | < 10E-10 |
| 3+ | 1,443 | < 10E-10 |

(f)

| mRNA feature | n | *p* |
|---|---|---|
| uORF number | 445 | <10E-6 |
| RNA structure (near cap) | 945 | <10E-6 |
| Kozak consensus | 161 | 0.01 |

**Fig. 2**

(c)

| Functional class | *p* |
|---|---|
| Transcription factors | <10E-10 |
| Transcription machinery | 1.0 |
| Messenger RNA biogenesis | 1.0 |
| Spliceosome | 1.0 |
| Cytoplasmic ribosome | <10E-10 |
| Mitochondrial ribosome | <10E-10 |
| Translation factors | <10E-5 |

| | |
|---|---|
| Chaperones and folding catalysts | <10E-6 |
| Membrane trafficking | 0.2 |
| Ubiquitin system | <10E-5 |
| Proteasome | <10E-10 |
| Glycolysis | <10E-6 |
| Pentose phosphate pathway | <10E-6 |
| TCA cycle | 0.002 |
| Fatty acid biosynthesis | 0.01 |
| Fatty acid degradation | 0.01 |
| Oxphos | <10E-5 |
| Nucleotide metabolism | <10E-4 |
| Amino acid metabolism | <10E-4 |

(d)

| Start position | Length | n | *p* |
|---|---|---|---|
| 1 | 0 | 6,883 | Reference |
| 1 | 1 | 1,835 | <10E-10 |
| 1 | 2 | 449 | <10E-10 |
| 1 | 3 | 178 | <10E-10 |
| 1 | 4 | 101 | <10E-10 |
| 1 | 5 | 53 | <10E-10 |
| 1 | 6 | 36 | <10E-10 |
| 1 | 7 | 22 | <10E-4 |
| 1 | 8+ | 32 | <10E-10 |
| 2 | 0 | 3,507 | Reference |
| 2 | 1 | 4,675 | 0.03 |
| 2 | 2 | 1,066 | <10E-10 |
| 2 | 3 | 240 | <10E-3 |
| 2 | 4 | 72 | 1.0 |
| 2 | 5 | 17 | 1.0 |
| 2 | 6 | 7 | 1.0 |
| 2 | 7 | 3 | 1.0 |
| 2 | 8+ | 2 | 1.0 |
| 3 | 0 | 6,731 | Reference |
| 3 | 1 | 2,488 | 1.0 |
| 3 | 2 | 276 | 1.0 |
| 3 | 3 | 64 | 1.0 |
| 3 | 4 | 22 | 1.0 |
| 3 | 5 | 4 | 1.0 |
| 3 | 6 | 3 | 1.0 |
| 3 | 7 | 1 | 1.0 |
| 3 | 8+ | 0 | NA |

(e)

| Treatment | uORF number | n | *p* |
|---|---|---|---|
| No treatment | 0 | 5,635 | Reference |
| No treatment | 1 | 2,037 | <10E-10 |
| No treatment | 2 | 948 | <10E-10 |
| No treatment | 3+ | 969 | <10E-10 |
| Torin 1 | 0 | 5,635 | Reference |
| Torin 1 | 1 | 2,037 | <10E-3 |
| Torin 1 | 2 | 948 | 0.04 |
| Torin 1 | 3+ | 969 | <10E-6 |

(f)

| Treatment | CDS length | n |
|---|---|---|
| No treatment | (100, 178] | 27 |
| No treatment | (178, 316] | 376 |
| No treatment | (316, 562] | 1,087 |
| No treatment | (562, 1000] | 2,461 |
| No treatment | (1000, 1778] | 3,143 |
| No treatment | (1778, 3162] | 1,791 |
| No treatment | (3162, 5623] | 556 |
| No treatment | (5623, 10000] | 126 |
| No treatment | (10000, 17783] | 22 |
| Torin 1 | (100, 178] | 27 |
| Torin 1 | (178, 316] | 376 |
| Torin 1 | (316, 562] | 1,087 |
| Torin 1 | (562, 1000] | 2,461 |
| Torin 1 | (1000, 1778] | 3,143 |
| Torin 1 | (1778, 3162] | 1,791 |
| Torin 1 | (3162, 5623] | 556 |
| Torin 1 | (5623, 10000] | 126 |
| Torin 1 | (10000, 17783] | 22 |

**Fig. 3-5 and Supplementary Fig. 4-8**

Mean ribosome load for each condition was calculated as the combined average of the biological replicate data:

n = 3 (RCC4, RCC4 VHL, 786-O and 786-VHL)

n = 2 (RCC4 with Torin 1, RCC4 VHL with Torin 1, 786-O (*EIF4E2* KO; one clone was generated using g1 gRNA and a second using g2 gRNA), and 786-O VHL (*EIF4E2* KO; one clone was generated using g1 gRNA and a second using g2 gRNA)

mRNA abundance for each condition was calculated as the combined average of the biological replicate data:
n = 3 (RCC4 and RCC4 VHL)
n = 4 (786-O and 786-O VHL)

**Fig. 3**
(e)

| Cell | TOP motif length | n | *p* |
|---|---|---|---|
| RCC4 | 0 | 8,206 | Reference |
| RCC4 | 1 | 2,211 | <10E-10 |
| RCC4 | 2 | 532 | <10E-10 |
| RCC4 | 3 | 213 | <10E-10 |
| RCC4 | 4 | 120 | <10E-10 |
| RCC4 | 5 | 63 | <10E-10 |
| RCC4 | 6 | 38 | <10E-10 |
| RCC4 | 7 | 29 | <10E-5 |
| RCC4 | 8+ | 35 | <10E-10 |
| 786-O | 0 | 5,902 | Reference |
| 786-O | 1 | 1,723 | 1.0 |
| 786-O | 2 | 374 | 1.0 |
| 786-O | 3 | 135 | 1.0 |
| 786-O | 4 | 105 | 1.0 |
| 786-O | 5 | 46 | 1.0 |
| 786-O | 6 | 30 | 1.0 |
| 786-O | 7 | 21 | 1.0 |
| 786-O | 8+ | 27 | 1.0 |

**Supplementary Fig. 3**
(a)
Significance of mRNA features to predict mean ribosome load

| Segment | RNA feature | n | *p* |
|---|---|---|---|
| 5' UTR | Length | 12,268 | <10E-10 |
| 5' UTR | uORF number | 12,268 | <10E-10 |
| 5' UTR | RNA structure (near cap) | 12,266 | <10E-10 |
| 5' UTR | RNA structure (distal) | 7,325 | <10E-6 |
| CDS | Kozak consensus | 12,268 | <10E-10 |
| CDS | Length | 12,268 | <10E-10 |
| CDS | RNA structure | 12,268 | <10E-10 |

Calculation of $R^2$ (only n for iteration 1 is shown as a representative data)

| Iteration | Segment | RNA feature | n (training) | n (test) |
|---|---|---|---|---|
| 1 | 5' UTR | Length | 9,469 | 1,457 |
| 1 | 5' UTR | uORF number | 9,469 | 1,457 |
| 1 | 5' UTR | RNA structure (near cap) | 9,468 | 1,457 |
| 1 | 5' UTR | RNA structure (distal) | 5,607 | 766 |
| 1 | CDS | Kozak consensus | 9,469 | 1,457 |
| 1 | CDS | Length | 9,469 | 1,457 |
| 1 | CDS | RNA structure | 9,469 | 1,457 |

(b)

| MFE (-kcal/mol/nt) | n | *p* |
|---|---|---|
| [0.0064, 0.278] | 2,454 | Reference |
| (0.278, 0.344] | 2,454 | Reference |
| (0.344, 0.402] | 2,450 | Reference |
| (0.402, 0.474] | 2,454 | Reference |
| (0.474, 1.1] | 2,454 | <10E-10 |

(d)

| Kozak consensus score | n | *p* |
|---|---|---|
| [0.164,0.56] | 2,454 | <10E-10 |
| (0.56,0.642] | 2,454 | 0.002 |
| (0.642,0.712] | 2,453 | Reference |
| (0.712,0.785] | 2,453 | 0.03 |
| (0.785,0.989] | 2,454 | 0.02 |

**Supplementary Fig. 4**
(b)

| CDS length | n | *p* |
|---|---|---|
| (100, 178] | 27 | <10E-4 |
| (178, 316] | 376 | <10E-10 |
| (316, 562] | 1,087 | <10E-3 |
| (562, 1000] | 2,461 | <10E-10 |
| (1000, 1778] | 3,143 | <10E-10 |
| (1778, 3162] | 1,791 | <10E-10 |
| (3162, 5623] | 556 | <10E-10 |
| (5623, 10000] | 126 | <10E-10 |
| (10000, 17783] | 22 | <10E-9 |
